# Peptide Sequence Features that Reduce Membrane Pore Line Tension

**DOI:** 10.64898/2026.09.06.749768

**Authors:** Denys Biriukov, Robert Vácha

**Affiliations:** CEITEC – Central European Institute of Technology, Masaryk University, Kamenice 753/5, 625 00 Brno, Czech Republic; National Centre for Biomolecular Research, Faculty of Science, Masaryk University, Kamenice 753/5, 625 00 Brno, Czech Republic; Department of Condensed Matter Physics, Faculty of Science, Masaryk University, Kotlářská 267/2, 611 37 Brno, Czech Republic

## Abstract

Membrane pore stability is central to many processes involving membrane permeabilization, yet it remains unclear how the sequence of pore-localizing peptides can modify the energetics of the pore boundary. For large pores, the energetic cost associated with increasing pore size is described by the membrane line tension. Peptides capable of reducing line tension can therefore stabilize permeable membrane states, making their identification relevant for the design of membrane-active molecules. Here, we combine coarse-grained molecular dynamics simulations, free energy calculations, and evolutionary optimization algorithms to identify sequence features of *α*-helical peptides that reduce membrane line tension. The best-performing peptides consistently showed an amphipathic organization with aromatic-rich termini, a hydrophobic/aromatic membrane-facing nonpolar face, and a negatively charged polar face. These sequence features promoted peptide localization at the pore rim close to the intact bilayer and orientation parallel to the membrane edge. This binding geometry reorganized lipids at the pore rim and efficiently reduced the exposure of their hydrophobic tails to water, thereby reducing the energetic cost of the pore boundary. The main sequence and mechanistic trends identified in coarse-grained simulations were reproduced in all-atom simulations. Together, these results link peptide sequence to pore-rim localization, lipid reorganization, and ultimately line-tension reduction, providing molecular design principles for *α*-helical peptides that stabilize permeable membrane states.

## Introduction

Lipid membranes form the compartmental boundaries of cells and organelles, separating aqueous environments and controlling exchange with their surroundings.^1^ This barrier helps maintain electrochemical gradients and distinct chemical environments across cellular membranes.^1^ Membrane permeability can increase when membranes are perturbed by external stimuli, such as an electric field,^2^ or membrane-active molecules, *e.g.*, antimicrobial peptides.^3,4^ In many cases, these perturbations increase permeability by inducing membrane defects or pores that allow molecular transport across the membrane.^5^ Understanding how these permeable states arise and persist at the molecular level is important for explaining membrane disruption and designing membrane-active compounds.

Membrane defects and pores span different sizes and structures, ranging from small local disruptions to larger water-filled channels.^6^ Small defects are governed by local lipid rearrangements, water penetration, and the exposure of hydrophobic lipid regions to the aqueous phase.^7^ For larger pores with a recognizable boundary, these molecular contributions are commonly conceptualized through line tension,^8,9^ which is defined as the free energy per unit length of the pore boundary. For a given membrane composition, line tension reflects the energetic cost to change the size of the pore. ^7,10^ Larger values increase the thermodynamic driving force for pore closure, whereas reducing this cost can prolong pore lifetime. ^7,10^ Molecules that localize at the pore boundary can alter the membrane line tension. ^11,12^ Such molecules can modulate pore stability by shielding unfavorable lipid exposure, changing local lipid organization, and thereby altering the free energy cost to open and increase the size of the pore.^9^

Amphipathic peptides are well suited to modulate pore energetics because their sequences can encode membrane binding, localization near the pore boundary, interactions with both the hydrophobic membrane interior and the aqueous phase, and local contacts with lipid headgroups.^5^ This combination of peptide features is central to the activity of many antimicrobial peptides, which often act by disrupting membrane integrity and causing leakage of cellular contents.^5^ In some cases, peptides have been suggested to form toroidal pores,^11–13^ in which the pore edge is covered by both peptides and lipid headgroups. For this type of pore, line tension provides a natural description of the energetic cost associated with changing the length of the pore boundary. This is different from, *e.g.*, barrel-stave pores, where peptides insert across the bilayer and assemble into well-defined channels with peptide-lined walls.^14,15^ Because toroidal pores involve lipids as part of the pore boundary, their stability depends not only on peptide assembly but also on how peptides reorganize lipids at this boundary. Toroidal pores are commonly heterogeneous and transient structures, making them difficult to characterize at the molecular level. ^16^ Indeed, even for several antimicrobial peptides for which toroidal pores have been suggested as a mechanism of action, the evidence is often indirect or inferred from theoretical models.^11–13,17–19^

Therefore, predicting which peptide sequences will stabilize membrane pores remains a major challenge. Peptide sequence space is enormous, and although general properties such as net charge, hydrophobicity, and amphipathicity may provide useful guidance, they do not guide how the identity and position of individual amino acids control peptide behavior in a given membrane context. As a result, it remains unclear which sequence features can reliably modulate pore energetics, limiting the rational design of peptides that stabilize permeable membrane states.

Here, we combine coarse-grained molecular dynamics (CG-MD) simulations, free energy calculations, and evolutionary algorithms to identify peptide sequence features that reduce the effective line tension of membrane pores. To make this screening feasible, we build on our previously developed collective variable (CV) for estimating peptide-induced free energy changes of the pore line tension. ^7^ Representative sequences identified in the CG-MD screening are further examined using all-atom molecular dynamics (AA-MD) simulations to assess their behavior at higher molecular resolution. Together, this multiscale approach provides a computational framework for identifying peptide features that stabilize toroidal pores in lipid membranes.

## Methods

### Molecular Dynamics Setup

We used the molecular dynamics (MD) setup introduced in our recent work,^7^ where it was applied together with the “Rapid” CV to estimate membrane line tension. In this approach, a lipid stripe is used to model a toroidal pore of effectively “infinite” size, Figure 1A. The lipid stripe was generated using a similar procedure as in our previous work. ^7^ Briefly, a lipid bilayer initially generated with CHARMM-GUI^20–22^ and subsequently equilibrated was placed in a simulation box extended along one lateral dimension, allowing lipids at the two exposed membrane edges to reorient and form the lipid stripe. The only modification introduced here was the placement of an equal number of peptides at each membrane edge during equilibration. Unless stated otherwise, four peptides were placed at each edge with the same termini orientation, parallel to one another, and with approximately uniform spacing. Peptide termini were kept neutral to focus on sequence-dependent effects and to avoid additional variability arising from orientation-dependent interactions of charged termini. This choice is also consistent with many experimental studies, in which at least one peptide terminus is uncharged via capping.^14,15^ Each peptide was restrained from moving into the bilayer region of the lipid stripe, while remaining free to desorb from the rim. The detailed preparation protocol for the lipid stripe/peptide systems is provided in the Supporting Information (SI). The lipid stripe contained 200 lipids with a symmetric leaflet composition of 1-palmitoyl-2-oleoyl-*sn*-glycero-3-phosphoethanolamine (POPE) and 1-palmitoyl-2-oleoyl-*sn*-glycero-3-phosphoglycerol (POPG) at a 3:1 ratio. This composition was used as a simplified model of the inner membrane of *E. coli*,^23,24^ motivated by the potential relevance of line-tension-reducing peptides as antimicrobial agents. The lipid stripe was solvated with 6000 polarizable water beads (see the Coarse-Grained Molecular Dynamics section). Counterions, sodium or chloride, were added to neutralize the charge of POPG lipids and, when required, charged peptides. In addition, 65 NaCl pairs were added to mimic physiological salt conditions.

**Figure 1:**
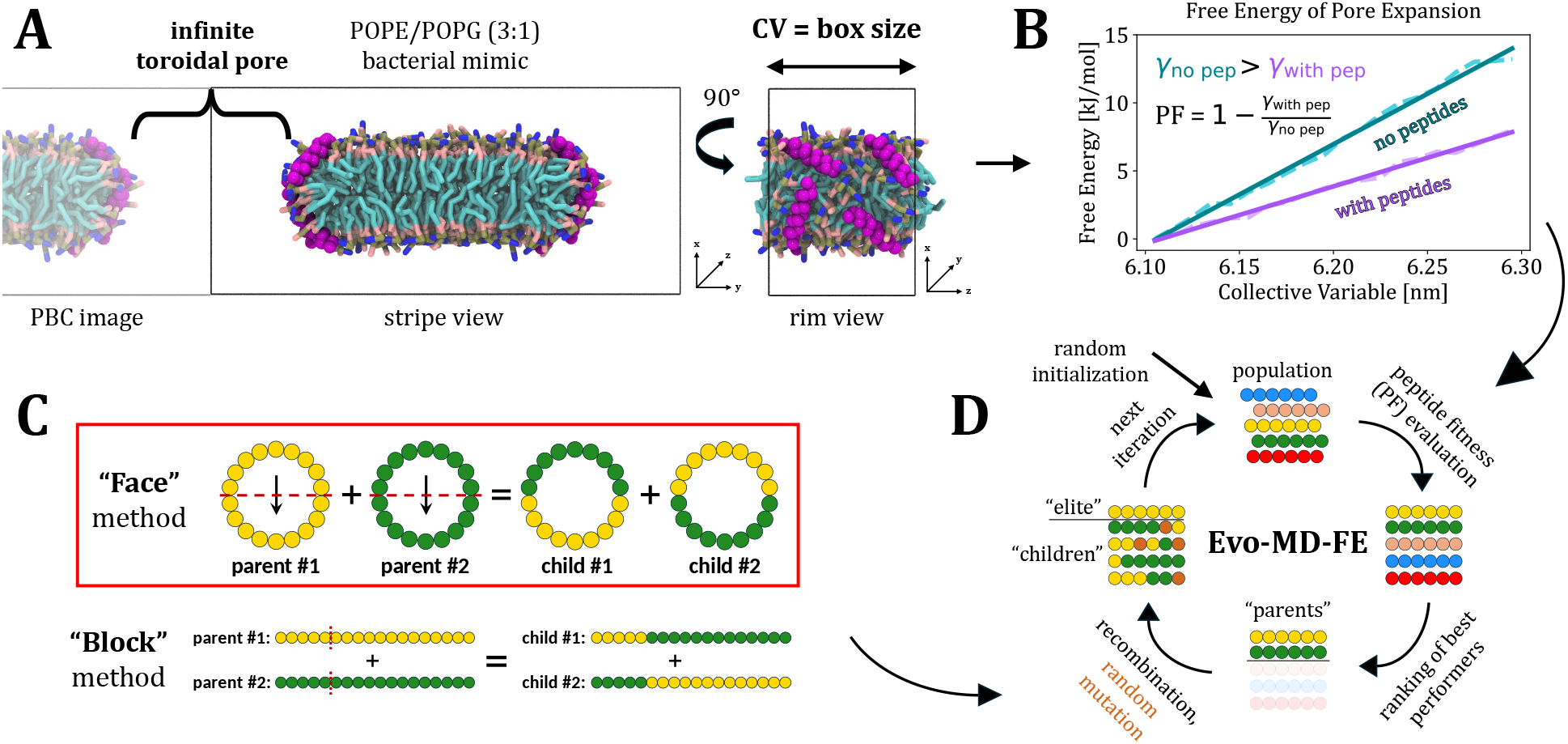
A–D) Methodology for identifying peptides that decrease line tension. **A) Molecular dynamics setup.** A lipid stripe with four identical peptides placed on each rim is used. The box size along the rim serves as a collective variable (CV) to evaluate the free energy of pore expansion; see Rapid method in Ref. 7 for details. **B) Peptide fitness estimation.** The free energy is fit with a linear function, whose slope corresponds to the line tension (*γ*). Peptide fitness (PF) is then calculated as the negative change in line tension relative to the peptide-free lipid stripe (PF *>* 0: decreased *γ*; PF *<* 0: increased *γ*). The genetic algorithm searches for peptides with the highest PF. **C) Recombination methods.** Two methods were used to generate “children” in the genetic algorithms. In the Face method, the helical wheels of the parent peptides are first aligned according to their hydrophobic moments. Each sequence is then divided into two equal halves across the helical wheel (nine residues per half for an 18-residue peptide), and the corresponding halves are exchanged between two parent peptides to generate two children. In the Block method, the primary sequence is split at a randomly chosen position, and the resulting segments are exchanged between two parent peptides. Most results presented in the main text were generated using Face method (highlighted in red rectangle). **D) Evolutionary algorithm workflow (Evo-MD-FE).** Initial peptides are generated randomly and evaluated using a peptide fitness function. The best-performing peptides are then selected as parents to produce offspring. Next, all peptides except the top performers (the “elite”) undergo random mutation, followed by another round of evaluation. This cycle is repeated iteratively.

### Free Energy Calculations

Free energy calculations were performed using the CV introduced in our previous work and referred to as Rapid.^7^ To evaluate line tension, we performed umbrella sampling (US) simulations using the box size along the pore rim as the CV. A series of simulations was performed at gradually varied box sizes, with the box size restrained in each window using the PLUMED action RESTRAINT. In our original work,^7^ we used 21 evenly spaced US windows covering pore-rim lengths from 6.0 to 6.6 nm, together with a force constant of 5000 kJ*·*mol*^−^*^1^*·*nm*^−^*^2^. Here, to reduce the computational cost, we used only three US windows, with equilibrium box sizes of 6.1, 6.2, and 6.3 nm. The narrower range of sampled box sizes also limits changes in the effective peptide coverage of the pore rim, which is important because the number of peptides is fixed while the rim length varies. A weaker restraint force constant of 1000 kJ*·*mol*^−^*^1^*·*nm*^−^*^2^ was applied along the CV, which provided sufficient sampling for estimating line tension, see Figure S2 in the SI. Free energy profiles were reconstructed using the gmx wham utility.^25^ For each CG-MD US window, a 300 ns production run was performed, and the final 200 ns was used for analysis. For each AA-MD US window, production runs were 250 ns long, or extended to 500 ns in selected cases, with the final 200 ns or 400 ns used for analysis, respectively. The same simulation lengths and analysis windows were used for peptide simulations performed outside the Evo-MD-FE optimization runs. Depending on the peptide and simulation level, each peptide was evaluated using one to four independent replicas to assess the consistency of the estimated line-tension reduction across repeated simulations.

The resulting free energy profiles were fitted with a linear model over the 6.1–6.3 nm range. The fitted slope was used to obtain the line tension, *γ*, according to *γ* = *m/*(2 *× N_A_*), where *m* is the fitted slope and the factor of two accounts for the presence of two pore rims. Representative free energy profiles obtained in the absence and presence of peptides are shown in Figure 1B. Additional supporting data for this free energy setup are provided in the SI.

### Evolutionary Algorithm

We used an evolutionary algorithm coupled to CG-MD simulations, referred to as “EvoMD”, which was previously introduced by Risselada’s group.^26,27^ EvoMD is an optimization method that integrates MD simulations into a genetic algorithm inspired by Darwinian evolution, making it well suited for identifying promising peptide candidates in large, discrete search spaces. In this approach, an initial population of candidate peptides is generated randomly, and each candidate is evaluated according to a predefined “peptide fitness” (PF) function. The best-performing candidates are selected as “parents” for the next generation, while genetic operations, such as crossover recombination and random point mutations, are applied to generate new peptides, referred to as “children”. A subset of the best-performing candidates, referred to as the “elite”, is carried over to the next generation unchanged to preserve high-PF solutions. The newly generated children and the retained elite peptides are then evaluated again using the PF function. Repeating this procedure over multiple generations progressively refines the peptide population toward improved PF. One of the key advantages of the EvoMD method is that it can rely on relatively short CG-MD simulations for fitness evaluation, because even if these simulations are not fully converged and do not provide accurate absolute observables, genetic-algorithm selection primarily depends on the relative ranking of candidates within a population, such that evolution can proceed in the correct direction as long as better-performing candidates outperform most alternatives.^28^ A schematic representation of the algorithm is shown in Figure 1D.

Here, we combined EvoMD with free energy calculations based on the Rapid CV and refer to this implementation as “Evo-MD-FE”. The PF was defined as the change in membrane line tension upon peptide insertion, Figure 1B. Because the objective was to identify peptides that decrease line tension, corresponding to PF *>* 0, the Evo-MD-FE algorithm was designed to favor peptides with higher PF values.

Another important aspect of our Evo-MD-FE method is the procedure used to generate children. In previous implementations of peptide evolutionary algorithms, ^26,27^ crossover recombination has typically been performed by splitting the primary sequences of two parent peptides into two segments and exchanging these segments between the parents. We refer to this sequence-based crossover as the “Block” method, Figure 1C. However, because all CG-MD simulations in this work were performed with peptides constrained in helical conformations, we mainly used an alternative crossover strategy, referred to as the “Face” method, Figure 1C, unless explicitly stated otherwise. In the Face approach, each parent peptide sequence is first mapped onto a helical wheel and aligned according to its hydrophobic moment. The hydrophobic moment was calculated using the amino-acid side chain hydrophobicity scale of Fauchère and Pliska. ^29^ The helical wheel is then divided into two faces of equal size along an axis perpendicular to the hydrophobic moment, and the corresponding faces are exchanged between two parent peptides. The helical-wheel representations of the newly generated peptides were then converted back into primary sequences; details of this procedure are provided in the SI. The use of the Face method was motivated by the expectation that effective line-tension-reducing peptides are likely to be amphipathic, with a relatively nonpolar face inserting into the hydrophobic interior of the lipid stripe and a more polar face oriented toward the aqueous solution and lipid headgroups. Thus, by operating directly on the amphipathic organization of helical peptides, the Face method was expected to guide the Evo-MD-FE search more efficiently toward candidates capable of stabilizing the membrane edge and lowering the line tension.

All Evo-MD-FE optimization runs used populations of 128 peptides per iteration. Each peptide was 18 residues long. At each iteration, the top-performing quarter of the population was selected as parents for generating children. Each residue in each generated peptide was then subjected to a random mutation with a probability of 1*/*18 (unless stated otherwise, see the SI), corresponding to one mutation per peptide on average. Two types of elite peptides were used, similar to previous EvoMD implementations.^26,27^ First, the two best-performing peptides from each iteration were passed unchanged to the next iteration; these peptides are referred to as “iteration elites”. Second, peptides that had already been evaluated in three independent runs were retained as “rerun elites”, with no more than two such peptides retained per iteration. Once a peptide had been evaluated three times, its existing PF value was reused in subsequent iterations without launching additional simulations.

Two amino acid sets were used for peptide generation. The first was a reduced amino acid set containing 10 residues (A, L, M, K, Q, E, W, S, Y, and F). This set was chosen to include residues with relatively higher *α*-helical propensity^30^ while retaining representatives of the major chemical classes. Using a reduced amino acid alphabet decreases the size of the sequence space to search and enables more efficient identification of sequence features associated with line-tension reduction. The second set was an expanded amino acid set containing 18 residues (A, L, M, K, Q, E, W, S, Y, F, G, V, D, R, H, T, C, and N). This set is based on the standard proteinogenic amino acids excluding proline (P), because it disrupts *α*-helical structure, and isoleucine (I), because its coarse-grained representation in the Martini force field used here is identical to that of leucine (L).^31^ Histidine (H) was modeled in its neutral form.

### Coarse-Grained Molecular Dynamics

CG-MD simulations were performed using a hybrid Martini 2.2 model developed in this work, referred to as M2.2hps. The model retains the Martini 2.2P representation of peptide amino acids^31^ and uses polarizable Martini water,^32^ together with rescaled peptide–peptide interactions,^33^ while retaining the standard Martini 2.2 interaction matrix^31^ for most remaining nonbonded interactions. The use of this hybrid force field was motivated by three reasons. First, the polarizable Martini water model^32^ was needed to accurately capture membrane line tension, as demonstrated in our previous work.^7^ Second, the latest Martini 3 model^34^ has been reported to capture peptide curvature sensing less accurately, ^35^ which is relevant here because the peptides localize at the curved pore rim. Third, rescaled Martini 2.2 interaction matrix was used to improve the description of peptide–peptide interactions, which are known to be overestimated in Martini 2 FF^33,36^ and which we also found to be problematic in its polarizable variant during test simulations. A detailed discussion of the force field choice, together with additional validation data supporting the use of this model, is provided in the SI.

The atomistic *α*-helical peptide structures, consisting only of heavy atoms, were first built using MODELLER,^37^ version 10.6. Martinize2 program^38^ was then used to generate the corresponding coarse-grained topologies with the Martini 2.2 polarizable mapping scheme for amino acids. The peptide secondary structure was constrained to remain helical in all CG-MD simulations, following the standard Martini force-field treatment of *α*-helical peptides. All production CG-MD simulations were performed with GROMACS^39^ (versions 2022.3, 2024.3, and 2025.4) in combination with PLUMED (versions 2.8.1, 2.9.3, and 2.9.4, respectively). We followed a standard protocol for Martini simulations.^40^ Periodic boundary conditions were applied in all directions, and neighbor searching was performed using the Verlet cutoff scheme with a neighbor-list cutoff of 1.35 nm.^41^ Reaction-field electrostatics and Lennard-Jones interactions were shifted to zero at a cutoff of 1.1 nm. For electrostatics, a relative dielectric constant of 2.5 was used within the cutoff, with an infinite dielectric constant beyond the cutoff. The membrane and solvent (peptides, ions, and water) groups were coupled separately to a stochastic velocity-rescaling thermostat^42^ at 310 K, using a coupling time constant of 1 ps. The pressure was maintained at 1 bar using an anisotropic Parrinello–Rahman barostat^43^ with a coupling time constant of 12 ps and a compressibility of 3*×*10*^−^*^4^ bar*^−^*^1^. As in our previous work, ^7^ the two membrane lateral box dimensions were coupled independently, *i.e.*, along the extended dimension and parallel to the pore rim. A time step of 20 fs was used in all production CG-MD simulations. Equilibration simulations used slightly modified protocols, which are described in detail in the SI.

### All-Atom Molecular Dynamics

AA-MD simulations were performed to validate the performance of peptides identified in the Evo-MD-FE optimization runs based on CG-MD simulations. For each selected peptide, equilibrated configurations taken from the end of the corresponding CG-MD US windows were backmapped to atomistic CHARMM-force-field-compatible systems using the CG2AT2 tool.^44^ As in the CG-MD simulations, peptide termini were modeled in their neutral form. Each configuration was equilibrated sequentially in the *NVT* and *NPT* ensembles, using a 1 fs time step for the 50 ps *NVT* equilibration and a 2 fs time step for the subsequent 1 ns *NPT* equilibration. The backmapped systems were then subjected to the same US protocol as used in the CG-MD simulations. Note that the *α*-helical conformation of the peptides was also maintained in AA-MD simulations by applying backbone dihedral restraints to the *ϕ* and *ψ* angles of each residue, using reference values of –60*^◦^* and –45*^◦^*, respectively, with a force constant of 500 kJ*·*mol*^−^*^1^*·*rad*^−^*^2^.

All AA-MD simulations were performed with GROMACS in combination with PLUMED, using the same versions as for the CG-MD simulations. We followed a standard protocol for CHARMM force field simulations. The CHARMM36^45^ was used for lipids, the CHARMM36m was used for peptides,^46^ the CHARMM-specific TIP3P model was used for water,^47,48^ and default CHARMM parameters were used for ions. The leap-frog integrator was used with a time step of 2 fs for all production simulations. Buffered Verlet lists^49^ were used for neighbor searching. Long-range electrostatics were treated using smooth particle-mesh Ewald (PME) algorithm.^50,51^ The input real-space cutoff was set to 1.2 nm, and PME tuning was enabled, allowing GROMACS to increase this value when required for optimal PME performance. Lennard-Jones interactions were cut-off at 1.2 nm, with forces smoothly switched to zero from 1.0 nm to 1.2 nm. The Nośe–Hoover thermostat^52,53^ was applied separately to lipids and to the non-lipid components, using a target temperature of 310 K and a coupling time constant of 1 ps. The Parrinello–Rahman barostat^43^ was applied anisotropically, as in the CG-MD simulations, using a reference pressure of 1 bar, a coupling time constant of 5 ps, and a compressibility of 4.5*×*10*^−^*^5^ bar*^−^*^1^. During equilibration, the Berendsen barostat^54^ with the same parameters was used instead. Water geometry was constrained using SETTLE,^55^ while all other covalent bonds involving hydrogen atoms were constrained using P-LINCS.^56,57^

### Simulation Analysis

#### Peptide orientation

Peptide orientation at the pore rim was quantified using the peptide end-to-end vector, defined between the first and last backbone beads of each peptide. For each peptide, the end-to-end vector was decomposed into its Cartesian components, and the absolute values of its normalized projections along the *y* and *z* axes were calculated as cos(dY) and cos(dZ), respectively. These values were averaged over all peptides present in the system, all analyzed frames, and all US windows for each peptide sequence.

#### Two-dimensional density profiles

Two-dimensional density maps were calculated separately for lipids and peptides. The membrane center of mass was translated to the origin in each frame to provide a common reference frame across the trajectories. Lipid distributions were calculated using either phosphate atoms or PO4 beads, whereas peptide distributions were calculated using either C*_α_* atoms or backbone beads. The distributions obtained from all three US windows were then combined, and two-dimensional histograms were constructed in the *xy* and *xz* planes using 1×1 Å^2^ bins, corresponding to views of the lipid stripe and pore rim, respectively. The resulting counts were normalized by the total number of trajectory frames, yielding the time-averaged two-dimensional number-density distributions shown in the density maps.

#### Radial distribution functions

Radial distribution functions (RDFs) between lipid tails and water were calculated using MDAnalysis.^58,59^ As for the two-dimensional density analysis, trajectories were first centered using the membrane center of mass to provide a consistent reference frame for defining the pore-rim regions. Lipids were assigned to the pore-rim regions in each trajectory frame according to the position of their PO4 bead, using two equivalent regions on opposite sides of the lipid stripe (x=–30 to 30 Å, y=–70 to –40 Å and x=–30 to 30 Å, y=40 to 70 Å). For lipids located within these regions, RDFs were calculated between the lipid-tail beads (C1A, D2A, C3A, C4A, C1B, C2B, C3B, and C4B) and uncharged water beads (W).

## Results and Discussion

### Sequence Features that Lower Pore Line Tension

To identify peptides that decrease pore line tension in the large-pore limit, we combined the Evo-MD-FE algorithm with CG-MD simulations and a novel free-energy CV to search for favorable sequence motifs. Because lower line tension reduces the energetic cost of pore expansion, these motifs are expected to define peptides capable of stabilizing toroidal membrane pores. In this work, we performed several Evo-MD-FE optimization runs under varying conditions, see Table S5 in the SI, most of which converged on the same overall trends and conclusions. In the main text, we focus on one representative Evo-MD-FE run, including subsequent AA-MD verification of the peptide sequences generated in this run, while the results from the additional runs are summarized in the SI and referred to where relevant in the main text. The main goal of this work was not to identify a single optimal peptide or determine the preferred amino acid at each sequence position, but rather to establish broader sequence features and design principles associated with line-tension reduction.

We performed an Evo-MD-FE optimization run using the Face method for generating new peptide sequences, as shown in Figure 1C. This optimization run used a reduced amino acid set (A, L, M, K, Q, E, W, S, Y, and F) and was carried out for 30 iterations. Excluding repeated evaluations of the best-performing peptides, this run tested 3,723 peptide sequences. As shown in Figure 2A, the optimization rapidly improved during the initial iterations and then showed a trend toward convergence, with the best-performing peptides in each iteration reducing the free-energy penalty associated with pore line tension by more than 40% (corresponding to PF *≈* 0.4). The average performance within each iteration remained at approximately 35%. Comparable performance was observed across the additional Evo-MD-FE runs, as summarized in the SI, Figures S7–S9. An additional run using the Block generation method, see Table S5, yielded slightly higher-performing peptides at the CG-MD level; however, subsequent AA-MD simulations showed that their performance was similar to that of the top peptides from the representative run discussed here.

**Figure 2:**
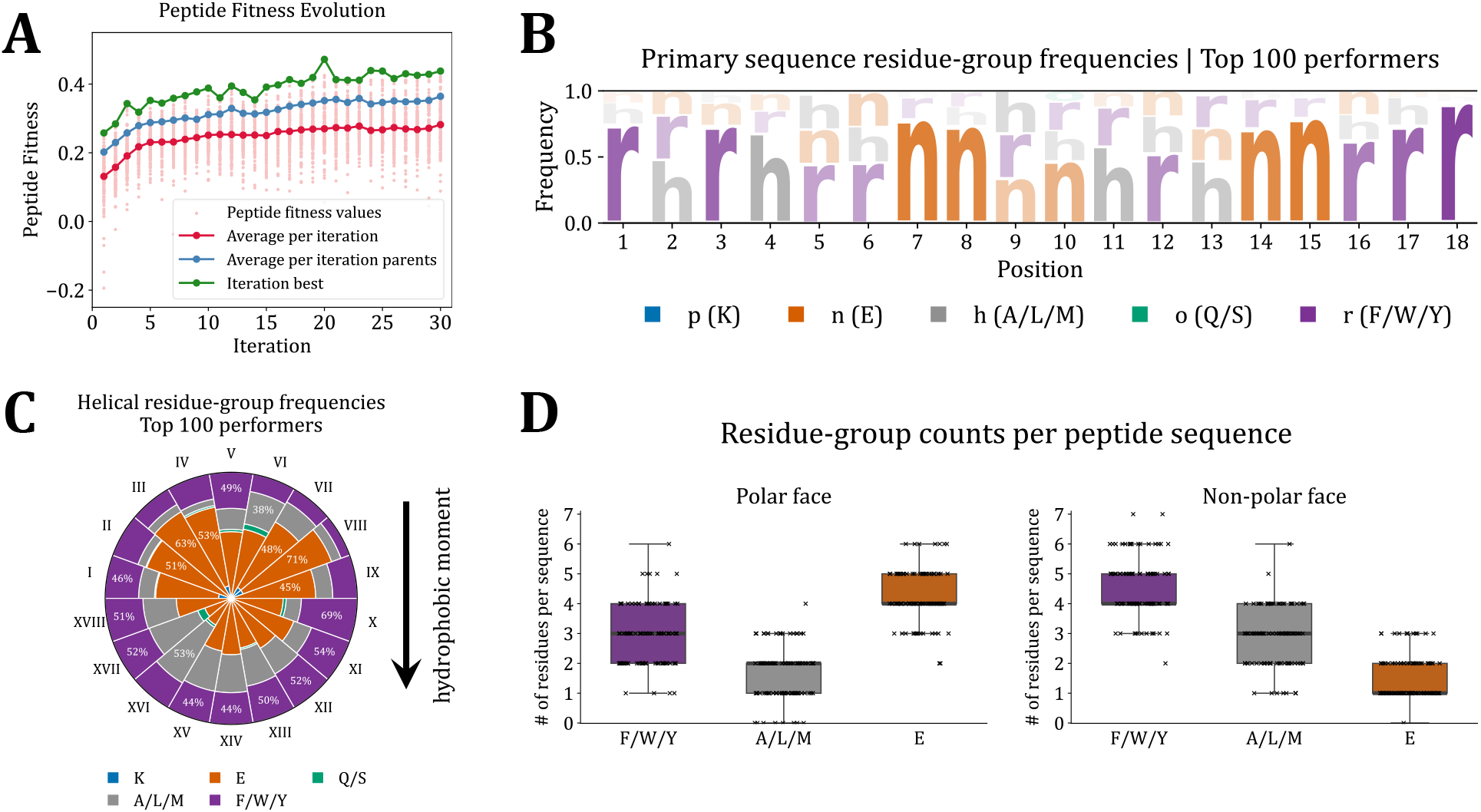
A–D) Evo-MD-FE peptide search with Face method of children generation. **A) Evolution of peptide fitness.** Peptide fitness (PF) values are shown as a function of iteration count. Each iteration contained 128 peptides, and only 10 amino acids (A, E, F, K, L, M, Q, S, W, Y) were used for peptide generation. For each iteration, all available PF values are reported (light-pink), either from a single simulation for a given peptide or as the average across all replicas available at that iteration. The plot also shows the mean PF per iteration (red), the mean PF of the parents (the best 32 peptides in each iteration; blue), and the best-performing peptide within each iteration (green). **B) Residue-group frequencies in the primary sequence.** Residues were grouped according to their physicochemical properties, namely positively charged, negatively charged, polar, hydrophobic, and aromatic. Consensus sequence logo for the top 100 peptides shows the positional enrichment of residue groups along the primary sequence. The letter height is proportional to the occupancy frequency at each sequence position. **C) Residue-group frequencies in the helical wheel.** Occupancy frequency of amino acid groups at each helical-wheel position among the top 100 peptides (top 100 peptides with the highest PF) is shown. Occupancies were calculated after aligning the helical wheel of each peptide with its hydrophobic moment vector. The area of each wheel segment is proportional to the relative frequency of the corresponding residue group at a given position. Positions I–IX correspond to the relatively more polar face, whereas positions X–XVIII correspond to the relatively more nonpolar face. Residue grouping is the same as in panel B. **D) Residue-group counts per peptide sequence.** Distributions of the numbers of aromatic residues (F/W/Y), hydrophobic residues (A/L/M), and negatively charged residues (E) located on the polar and nonpolar faces of the peptide helical wheel. Boxes indicate the interquartile range, thick horizontal black lines indicate the median, whiskers show the data range excluding outliers, and crosses represent individual peptide sequences.

We examined how peptide sequences evolved during this optimization and which sequence features were associated with the most effective line-tension reduction. For this analysis, we selected the top 100 performers from all peptides tested in this Evo-MD-FE run and analyzed their sequences in two complementary ways. First, the occurrence frequency of each amino acid was calculated at each position along the primary sequence. Second, each sequence was mapped onto a helical wheel, which was rotated according to its hydrophobic moment (again calculated using the hydrophobicity scale of Fauchère and Pliska^29^), after which the same occurrence frequencies were calculated at each wheel position (I–XVIII). In both cases, amino acids were then grouped according to their physicochemical properties. The resulting residue-group frequencies are shown in Figures 2B and 2C. We present the residue-group frequencies because they capture the dominant physicochemical trends without focusing too much on individual amino acids that may become enriched during Evo-MD-FE convergence. This representation is also consistent with the CG-MD resolution of the simulations (as well as with using the reduced amino acid set), where broader residue properties are expected to be more robustly resolved than fine residue-specific preferences. The corresponding individual amino-acid frequencies are nevertheless provided in the SI, Figure S4. Some specific amino-acid effects were also examined in targeted analyses discussed below.

The primary-sequence residue-group frequencies revealed a clear pattern among the top-performing peptides, Figure 2B. Aromatic residues were strongly enriched near the sequence termini, typically within the first three residues from either end, while a smaller fraction of nonpolar hydrophobic residues was also tolerated at these positions. In contrast, negatively charged residues were enriched closer to the middle of the sequence or, at least, depleted from the terminal regions.

Mapping the same sequences onto helical wheels revealed how these primary-sequence preferences translate into an amphipathic organization, Figure 2C. Aromatic residues were strongly enriched on one side of the wheel (positions X–XVIII), corresponding to the more nonpolar, membrane-facing face. This region also contained a substantial fraction of hydrophobic residues, namely alanine, leucine, or methionine. The opposite side (positions I–IX) was enriched in negatively charged glutamic acid residues, defining a pronounced, more polar and negatively charged face. Notably, uncharged polar residues (glutamine and serine) and positively charged lysine were almost entirely absent among the top performers. Together, these trends indicate that strong line-tension reduction is associated with aromatic-rich ends and amphipathic organization of the helix, combining a hydrophobic/aromatic face with a negatively charged polar face.

This amphipathic segregation was not strict, however, as aromatic and negatively charged residues were also present outside their dominant regions. In part, this reflects the primary-sequence constraints described above, since preferred residue positions along the sequence can map to slightly different regions after alignment of the helical wheels according to their hydrophobic moments. The diffuse pattern may also partly arise from the Face generation method, in which recombination between well-performing parent sequences can preserve favorable overall sequence features while introducing local deviations in the helical-wheel arrangement. Such variants can retain strong line-tension-reducing activity, making the position-specific amphipathic pattern less sharply defined across the full set of top performers. This interpretation is supported by the analysis of the top 20 performers (Figure S5 in the SI), for which the helical-wheel pattern is more sharply defined, indicating stronger convergence toward the preferred residue arrangement among the best line-tension reducers.

To complement the position-based analysis, we next examined the per-sequence residue composition of the top-performing peptides. Figure 2D shows the number of residues from each group located on the more polar and more nonpolar faces of the helical wheel. Overall, the best performers typically contained 6–9 aromatic residues, 3–6 hydrophobic residues, and 5–7 negatively charged residues. While the more nonpolar face typically tolerated one or two negatively charged residues, the more polar face frequently contained up to four aromatic residues. The presence of negatively charged residues on the more nonpolar face may provide additional polarity at the membrane-facing side, which could prevent overly deep insertion into the hydrophobic core and help maintain peptide localization near the pore rim. These negatively charged residues may also be positioned closer to the boundary between the polar and nonpolar faces, allowing the peptide to retain a sufficiently large hydrophobic patch. Conversely, the presence of aromatic residues on the more polar face likely reflects the strong preference for aromatic residues near the sequence termini, which translates into having some of these residues on the polar side of the helical wheel. This interpretation is supported by the corresponding helical-wheel analysis in which the first and last three residues of each sequence were excluded from the frequency calculation, substantially reducing the occurrence frequency of aromatic residues on the polar face, see Figure S6 in the SI. This suggests that aromatic residues on the polar face may not be required *per se*; rather, line-tension reduction may primarily depend on placing aromatic residues near the sequence termini while maintaining their preferential localization on the more nonpolar face whenever possible.

### Molecular Mechanism of Action to Reduce Membrane Pore Line Tension

A natural question arising from the observed sequence patterns is how these sequence features lead to the efficient reduction of membrane pore line tension. The answer appears to lie in the orientation and localization of the peptides at the pore rim and their effect on local rim lipid organization. Figure 3A shows the orientations of all peptides tested in the Evo-MD-FE optimization run described above. The best-performing peptides preferentially orient along the lateral membrane direction within the pore region and localize at the pore rim, relatively closer to the intact bilayer, see the schematics on the right side of Figure 3A. Similar orientation patterns were observed in the other Evo-MD-FE runs, as shown in Figure S10.

**Figure 3:**
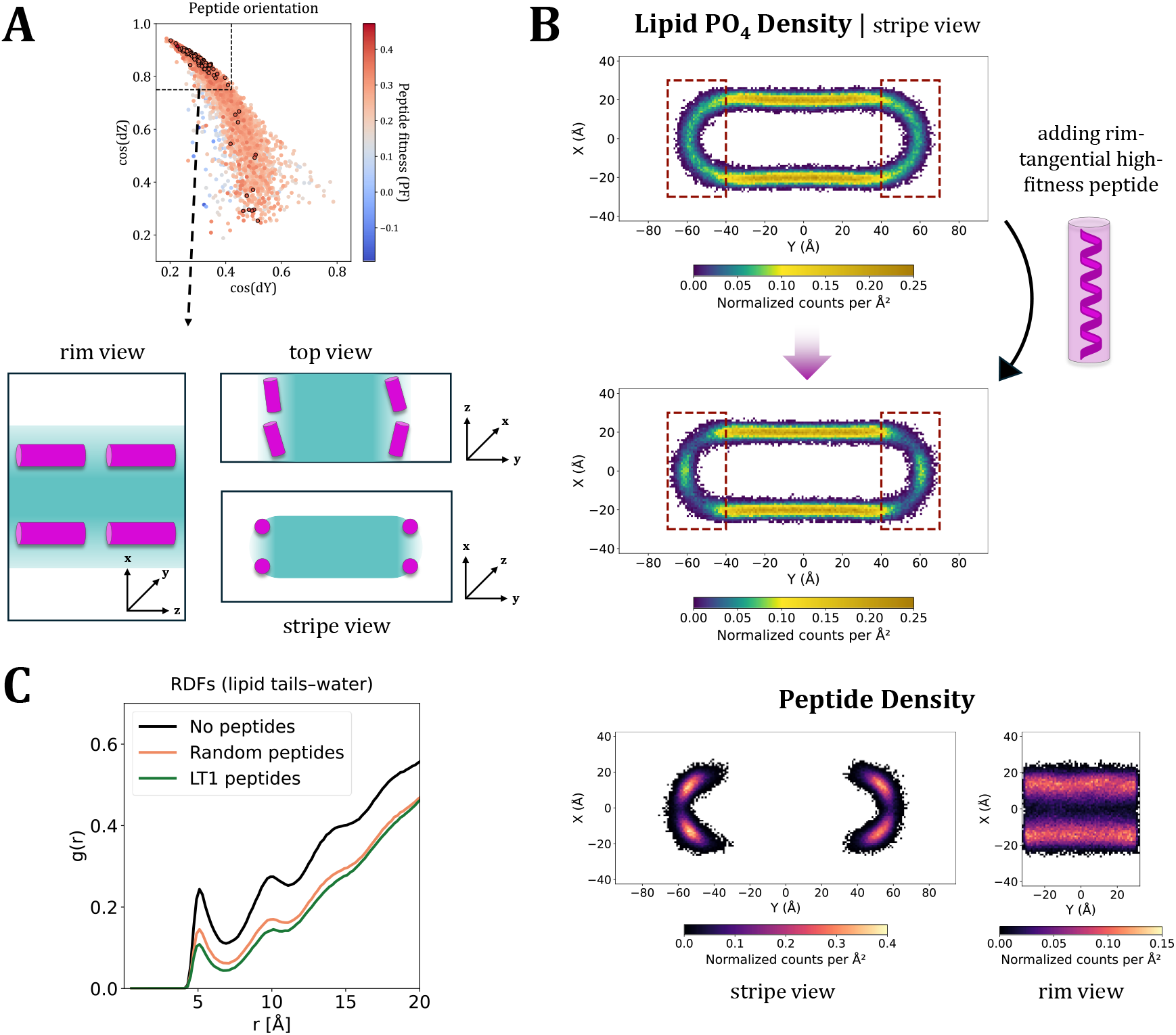
A–C) Mechanism of action for line tension-decreasing peptides. **A) Peptide orientation at the pore rim.** Orientation was evaluated for all peptides tested in the Evo-MD-FE run using the Face method. The orientation of each peptide was described by the absolute values of the normalized projections of the end-to-end vector connecting the first and last peptide backbone beads onto the *y* and *z* axes, reported as cos(*dY*) and cos(*dZ*), respectively. Calculated values represent averages across all eight peptides present in the system. Points are colored according to the corresponding peptide fitness (PF) value, and the top 100 performers are highlighted with a black outline. Each data point corresponds to a single simulation replica; values were not averaged across independent replicas. Simplified schematics on the right show the characteristic orientation adopted by the best performers, viewed along three coordinates. Pink cylinders represent peptides, while cyan regions represent lipids. Dashed lines are shown as guides to the eye. **B) Two-dimensional density maps of the stripe cross-section.** Lipid phosphate-group density maps are shown for both the peptide-free system (top) and the system containing high-fitness LT1 peptides (middle; see the main text). Dashed red rectangles highlight the rim regions. Peptide density maps for the same system (bottom) are shown in both stripe and rim views, illustrating preferential localization of the peptides at the pore rim. **C) Radial distribution functions (RDFs) between lipid tails and water.** Only lipids whose phosphate group was located within the highlighted regions shown in panel B in a given frame were included. RDFs are compared across three systems: the peptide-free system, a system containing random-sequence peptides from the Evo-MD-FE run that exhibits neither a high PF nor pronounced orientational behavior (panel A), and a system containing high-fitness LT1 peptides (see the main text).

Our interpretation of this behavior in relation to the preferred sequence features is as follows. The preferred orientation is likely promoted by the enrichment of aromatic residues at the sequence termini, which can anchor both peptide ends and align the peptide along the pore rim. This arrangement is likely favorable because it allows both ends of the helical peptide to remain associated with the membrane, rather than forcing the peptide to follow the curvature of the pore rim, which could weaken membrane contact at one or both ends. At the same time, negatively charged residues mainly located on the relatively more polar side of the peptide helical wheel may push lipid phosphate groups away from the peptide-rich parts of the rim through electrostatic repulsion. As a result, lipid headgroups are redistributed either toward the intact bilayer or toward the central part of the pore rim. This increases lipid headgroup density in these regions and makes the corresponding part of the rim more similar to an intact bilayer, with lipids more compactly packed and less unfavorable exposure of lipid tails to the polar environment. Such more bilayer-like organization may reduce the energetic cost of the pore rim, *i.e.*, the line tension. Our interpretation is supported by the two-dimensional density profiles shown in Figure 3B, which reveal lipid headgroup redistribution near the peptide-rich rim regions. The peptide density profiles further agree with the schematic orientation shown in Figure 3A. In addition, the localization of peptides at the rim, together with the presence of bulky aromatic residues, allows them to shield lipid tails from the aqueous environment, Figure 3C. Note that this shielding effect is present whenever any peptides bind at the rim, but it is stronger for the top-performing peptides.

To further test the proposed role of aromatic and negatively charged residues, we selected the best-performing sequence from the Evo-MD-FE optimization run, hereafter referred to as LT1, and introduced several controlled modifications to its sequence arrangement. First, we tested whether the primary-sequence placement of aromatic residues is indeed important for line-tension reduction. To do this, we generated 17 shifted variants of LT1, corresponding to shifts of one to seventeen residues relative to the original sequence, Figure 4A. These variants had the same amino-acid composition as LT1 and preserved the same relative residue arrangement on the helical wheel, but differed in the positions of residues along the primary sequence. This allowed us to isolate the effect of primary-sequence residue placement from the sequence composition and helical-wheel pattern. LT1 contained five aromatic residues within the terminal regions, defined here as the first and last three residues of the sequence, whereas all shifted variants contained only two to four terminal aromatic residues. Notably, all shifted variants performed worse than LT1, indicating that terminal enrichment of aromatic residues, and the peptide orientation and localization it promotes, is important for optimal line-tension reduction even when amino-acid composition and helical-wheel pattern are preserved.

**Figure 4:**
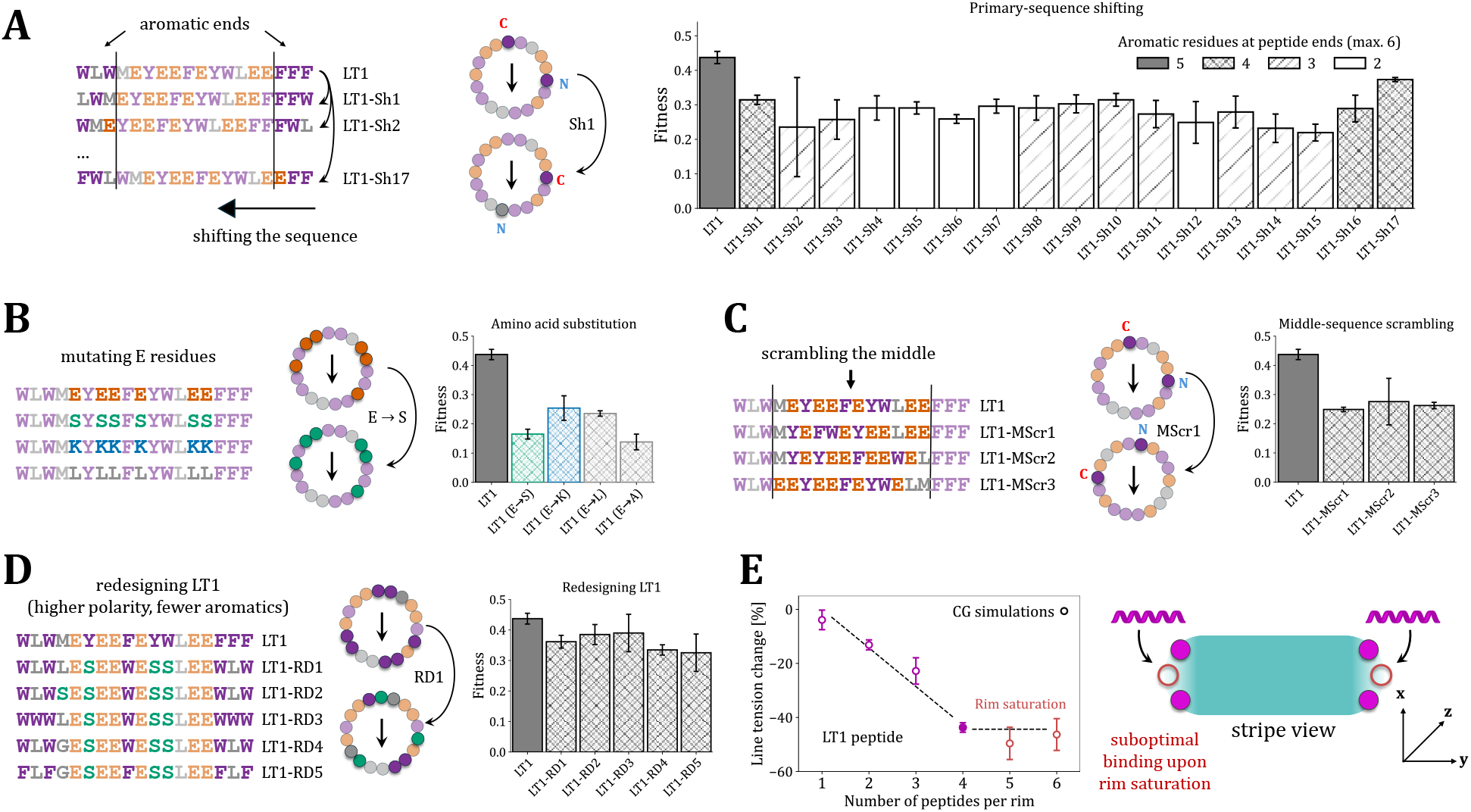
A–E) Sequence determinants and peptide-number dependence of line-tension reduction. **A) Primary-sequence shifting.** The selected top-performing peptide (LT1) was cyclically shifted to generate 17 variants with identical amino-acid composition and the same relative residue arrangement on the helical wheel, but different residue positions along the primary sequence. The calculated fitness values of LT1 and the shifted variants are compared. Bar shading indicates the number of aromatic residues (F/W/Y) within the peptide termini, defined as the first and last three residues. **B) Substitution of negatively charged residues.** All glutamate (E) residues in LT1 were replaced with serine (S), lysine (K), or leucine (L), while the remainder of the sequence was retained. The fitness values of LT1 and the resulting variants are compared. **C) Middle-sequence scrambling.** The aromatic-rich terminal regions of LT1 were retained, whereas the non-terminal residues were randomly shuffled while preserving their amino-acid composition. The fitness values of LT1 and the resulting variants are compared. **D) Rational redesign of LT1.** LT1 was redesigned to reduce aromatic content and increase polarity while retaining favorable line-tension-reducing features. The fitness values of LT1 and the redesigned variants are compared. **E) Dependence of line-tension reduction on the number of peptides at the membrane rim.** The relative line-tension change is shown for systems containing one to six LT1 peptides per rim. Increasing the number of peptides from one to four per rim progressively reduced the line tension, whereas adding further peptides produced no consistent additional reduction, due to the saturation of favorable binding positions at the rim. The schematic on the right illustrates the suboptimal binding of additional peptides after rim saturation. Dashed lines are guides to the eye. All error bars represent the standard deviation across replicate simulations.

We performed additional tests in which all terminal aromatic residues were replaced with either L or A, or the terminal positions were made fully aromatic using W, F, or Y, see Figure S11 in the SI. Replacing terminal aromatic residues with L or A strongly reduced peptide fitness. In the A-substituted variant, the rim-aligned orientation was altered, whereas the L-substituted variant largely preserved this orientation but showed a weaker shielding effect after replacement of the bulky aromatic residues by leucine. This suggests that hydrophobic residues can support the preferred orientation, whereas aromatic residues are additionally needed for stronger anchoring and more effective shielding of lipid tails. In contrast, fully aromatic ends generally retained high fitness, as well as the preferred peptide orientation and localization, but replacing the second-position L in LT1 with an aromatic residue did not further improve peptide performance. This indicates that fully aromatic ends are not necessarily the most optimal configuration. Among the aromatic residues, W generally performed best, followed by F and Y. Across the Evo-MD-FE runs, the frequent enrichment of W and F supports the preference for these aromatic residues, while the consistent presence of non-aromatic hydrophobic residues indicates that mixed aromatic/hydrophobic segments are favored, both near the peptide termini and on the more nonpolar face of the helical wheel, rather than fully aromatic regions.

Next, we replaced all negatively charged residues (E) in LT1 with either polar residues (S), positively charged residues (K), or hydrophobic residues (L or A), Figure 4B. In all cases, the mutated peptides reduced the line tension less effectively. For the E*→*S and E*→*K variants, peptide orientation remained similar to that of LT1, Figure S12. In contrast, the E*→*L and E*→*A variants showed altered peptide orientation, indicating that aromatic-rich ends are not sufficient to determine the preferred rim-associated orientation. For example, additional hydrophobic residues may perturb this orientation, whereas negatively charged residues may help stabilize it through the preferred reorganization of lipids and peptides at the pore rim. Overall, our data indicate that the presence of the negatively charged residues is essential for strong line-tension reduction and that the orientation promoted by aromatic-rich ends is not sufficient on its own.

Next, we tested several LT1 variants with unchanged aromatic-rich ends but randomly shuffled non-terminal residues, Figure 4C. This shuffled the positions of all non-terminal residues on the helical wheel, including the negatively charged residues, which were no longer concentrated mainly on the more polar face. As a result, the overall amphipathic pattern of the peptide was altered. This also lowered the line-tension-reducing effect, indicating that aromatic-rich ends and negatively charged residues are not sufficient by themselves for efficient line-tension reduction. Rather, the overall placement of residues is important. Moreover, peptide orientation was notably altered, Figure S12 in the SI, indicating that aromatic-rich (or at least hydrophobic) ends are necessary but not sufficient to produce the preferred rim-associated orientation; a sufficiently large nonpolar residue patch on the more nonpolar face also appears to be required for favorable peptide binding. Therefore, efficient line-tension reduction depends on balanced amphipathic organization, combining aromatic-rich ends with negatively charged residues defining the polar face and hydrophobic/aromatic residues on the opposite face promoting the preferred binding arrangement at the pore rim. Using these molecular determinants, we next attempted to redesign LT1 to retain the features associated with strong line-tension reduction while reducing its overall aromatic content and increasing the number of polar residues. This was motivated by the high aromatic and hydrophobic content of the Evo-MD-FE-derived peptides and the possibility that a more polar composition could improve properties such as solubility. As shown in Figure 4D, the redesigned peptides retained near-LT1 performance despite containing as few as five aromatic residues (W or F) and up to four polar residues (S), in addition to negatively charged glutamic acid residues (E). Up to three aromatic residues were placed near each terminus, while an additional one was positioned in the middle of the sequence so that it occupied the more nonpolar face of the helical wheel. The peptides also retained a relatively large hydrophobic binding patch on this face. Four redesigned peptides used WLW or FLF ends, whereas one had WWW ends, allowing us to test whether partial replacement of terminal aromatic residues by leucine was sufficient to preserve the preferred peptide orientation. Overall, these redesigned peptides retained good line-tension-reducing capacity, while containing fewer aromatic residues and a more polar sequence composition.

Note that in all tests shown in Figure 4, PF remained at approximately 0.15 or higher, because even the mutated variants could still partly shield lipid tails from the polar environment upon binding and therefore retained some line-tension-reducing capacity. Thus, the key distinction is not whether a peptide has any effect, but whether that effect is large enough to meaningfully lower the energetic cost of pore-size changes.

Finally, we examined how the line-tension-reducing effect depends on the number of peptides present at the pore rim, Figure 4E. Because all Evo-MD-FE calculations were performed with four peptides per rim, we tested additional systems containing from one to six LT1 peptides per rim. The results indicate that increasing the peptide number from one to four progressively decreased the line tension. However, adding more LT1 peptides did not lead to a further consistent decrease. This likely reflects saturation of the preferred peptide-binding sites at the rim: once four peptides are present, the optimal binding positions are already occupied, and additional peptides cannot bind in the same favorable arrangement. Consistent with this interpretation, partial peptide unbinding was observed in some simulations containing five or six peptides per rim. However, four peptides per rim should not be interpreted as a general saturation number, but as the preferred occupancy for the pore size used here. For larger pores, the increased rim length would create additional favorable binding sites, allowing more peptides to bind in the optimal rim-associated orientation.

### Atomistic-Simulation Validation of Top Performers

All Evo-MD-FE optimizations, as well as additional verification simulations, were performed with the coarse-grained Martini model, because AA-MD simulations at this scale are not feasible. Nevertheless, it is essential to verify the main outcomes using AA-MD simulations. To this end, we selected several groups of peptides from the Evo-MD-FE optimizations, together with additional model peptides, to test specific trends observed in CG-MD. These groups included: (i) ten top performers (including LT1) from the Evo-MD-FE run described in the main text, which follow the discussed structural patterns and mechanism of action proposed in Figure 3; (ii) ten poor performers from the same optimization run (as a negative control), which mostly originated from early iterations because the optimization rapidly moved away from poorly performing regions of sequence space; (iii) two mediocre performers enriched in lysine residues, included to test whether positively charged residues might be systematically misrepresented by the CG-MD simulations in the context of this work; (iv) five top performers from another Evo-MD-FE run (run #4, see Table S5 in the SI), which also contain aromatic-rich sequence ends and follow the proposed mechanism, thereby increasing the sequence diversity of the tested high-performing peptides; (v) five symmetric model peptides that differ only in the identity of charged or polar residues (E, D, K, R, or S), designed to test whether line-tension-reducing peptides preferentially contain negatively charged residues rather than positively charged or polar residues; (vi) two simple amphipathic peptides representative of negative-and positive-curvature sensing peptides,^35^ included to test whether curvature sensing alone is sufficient to induce strong line-tension reduction; (vii) some of the selected LT1 derivatives tested in Figure 4; (viii) two examples of outliers emerged during the optimization procedure that either did not follow the sequence features or mechanism of action proposed in Figure 3, or showed poor agreement between CG-MD and AA-MD simulations.

The results shown in Figure 5 indicate overall good agreement between the CG-MD and AA-MD simulations. The high-performing peptides cluster in the same region in both representations, with a line-tension reduction of approximately 40% (corresponding to PF *∼* 0.4). LT1 was the best-performing peptide in the AA-MD simulations (except the outlier, see below), which is consistent with its performance in the CG-MD Evo-MD-FE run. Most poor CG-MD performers also showed lower efficiency in AA-MD simulations. However, these peptides did not form a compact cluster, most likely because they represent diverse sequence patterns that may be differently described at atomistic resolution.

**Figure 5:**
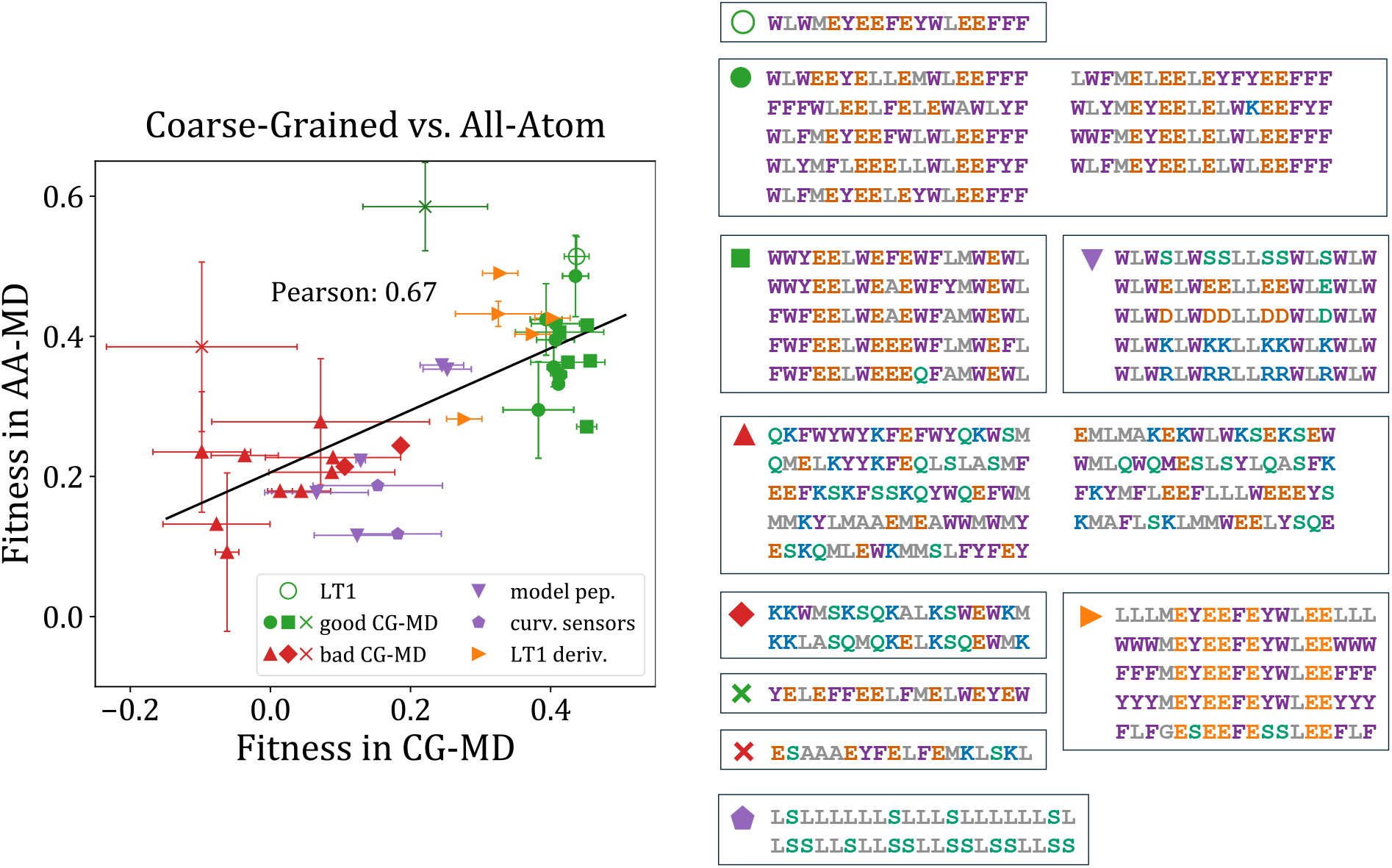
Comparison of peptide fitness from CG-MD and AA-MD simulations. Green symbols indicate peptides with favorable CG-MD performance, red symbols indicate peptides with unfavorable CG-MD performance, purple symbols indicate model peptides, and orange symbols indicate LT1-derived sequences. Different marker shapes distinguish the peptide subsets described in the main text: (i) circles, (ii) upward-pointing triangles, (iii) diamonds; (iv) squares, (v) downward-pointing triangles, (vi) pentagons, (vii) right-pointing triangles, and (viii) crosses. Horizontal and vertical error bars, where shown, represent standard deviations calculated from replicate CG-MD and AA-MD simulations, respectively. The black line shows the linear regression, with the corresponding Pearson correlation coefficient displayed on the graph. The sequences of all compared peptides are shown to the right of the graph.

The model peptides showed similar relative performance in the CG-MD and AA-MD simulations, further supporting the conclusions drawn throughout this work. The simple curvature-sensing peptides did not show notable line-tension-reducing activity at either simulation resolution, indicating that curvature sensing alone is not sufficient to explain the line tension reduction. The LT1-derived peptides generally performed worse than LT1, consistent with the CG-MD results, while the variants that retained high performance in CG-MD remained effective at the AA-MD level as well.

A few outliers also showed performance comparable to some of the best CG-MD performers. This is not entirely unexpected, because AA-MD simulations can resolve sequence-specific interactions that are not fully captured at CG-MD level. In addition, some peptides may reduce line tension through alternative mechanisms that are not represented by the main sequence patterns identified from the Evo-MD-FE optimization runs at the CG-MD resolution. For example, one outlier (green cross in Figure 5) has a sequence composition similar to that of the high-performing peptides discussed above, with multiple aromatic, negatively charged, and hydrophobic residues, but does not follow the same sequence template or preferred rim-associated arrangement. Instead, the peptides were more homogeneously distributed along the pore rim, Figure S13 in the SI. A second outlier (red cross in Figure 5) has a more distinct sequence composition, with aromatic residues concentrated near the middle of the sequence. These residues mediate binding near the tip of the pore rim, while the peptides overall displace lipid headgroups away from this region, Figure S13 in the SI. Interestingly, this peptide binding increases line tension in the CG-MD simulations but decreases it at the AA-MD level. For both outliers, however, the uncertainties are large, and partial unbinding events were observed in both CG-MD and AA-MD simulations. It should also be noted that AA-MD simulations are more difficult to converge because of the rougher atomistic energy landscape, so incomplete sampling may contribute to the observed differences. Nevertheless, these examples suggest that alternative line-tension-reducing mechanisms likely exist beyond the sequence and binding patterns identified here and could be explored in future work.

## Conclusions

In this work, we combined CG-MD simulations, free energy calculations, and evolutionary optimization to identify sequence features of *α*-helical peptides that reduce membrane pore line tension. Across independent optimization runs, the best-performing peptides converged toward a common set of features, namely aromatic-rich termini and an amphipathic organization with a hydrophobic/aromatic membrane-facing region and a negatively charged polar face. These features promoted peptide localization at the pore rim near the intact bilayer and orientation parallel to the membrane edge, where the peptides reorganized nearby lipids and efficiently shielded hydrophobic lipid tails from water. Targeted sequence modifications showed that these properties depend not only on overall composition but also on the precise placement of individual residues. Even among sequences that broadly satisfy the identified sequence features, relatively small changes can substantially reduce performance. The main sequence and mechanistic trends obtained from CG-MD simulations were also reproduced at the AA-MD level, supporting the molecular design principles identified here.

Several aspects remain to be addressed before these principles can be translated into experimentally testable peptides. Our simulations here characterize peptides already bound to the pore rim: peptides were initially placed close to the membrane edge (although they were free to desorb), and the free energy of membrane recruitment and pore targeting was therefore not considered explicitly. In particular, whether the negatively charged peptides identified here can efficiently target negatively charged membranes, despite favorable interactions of their aromatic residues with the membrane, remains to be established. Peptide solubility is another important consideration, particularly given the high aromatic content of many top-performing sequences. Conformational stability also remains to be checked, as the peptides were constrained/restrained in an *α*-helical conformation in both CG-MD and AA-MD simulations. Finally, because the present study does not address the initial formation of membrane pores, experimental validation may be most direct in systems where pores are generated independently, allowing peptide-mediated pore stabilization to be assessed separately from pore formation itself. Future work can therefore extend the present optimization toward membrane recruitment, solubility, and structural stability, while the sequence trends identified here can guide the selection of candidates for experimental validation and the design of membrane-active peptides with controlled effects on membrane permeability.

## Data Availability

The code and files required to reproduce the simulation workflows used in this work will be made available on Zenodo following peer review.

## Acknowledgement

This work was supported by the Czech Science Foundation (grant 26-20435S), the European Research Council (ERC) under the European Union’s Horizon 2020 research and innovation programme (grant agreement No 101001470), and the project National Institute of virology and bacteriology (Programme EXCELES, ID Project No. LX22NPO5103) - Funded by the European Union - Next Generation EU. We acknowledge VSB – Technical University of Ostrava, IT4Innovations National Supercomputing Center, Czech Republic, for awarding this project access to the LUMI supercomputer, owned by the EuroHPC Joint Undertaking, hosted by CSC (Finland) and the LUMI consortium through the Ministry of Education, Youth and Sports of the Czech Republic through the e-INFRA CZ (grant ID: 90254), project OPEN-35-3. The authors acknowledge the use of ChatGPT for assistance in improving the readability and language of the manuscript. The authors remain solely responsible for the scientific content and interpretations presented.

## Supporting Information Available

Additional methodological details on the molecular dynamics simulations with Rapid CV; description of the Martini hybrid force field; details of the additional Evo-MD-FE runs and the results derived from them; and supplementary data supporting the molecular determinants of the line-tension reduction described in the main text.

