## Supporting Information for "Peptide Sequence Features that Reduce Membrane Pore Line Tension"

### Molecular Dynamics Setup

#### CG-MD lipid stripe with added peptides

To prepare peptide-containing lipid stripes suitable for line-tension evaluation using the Rapid CV, we followed a protocol similar to that established in our original peptide-free study.[S1] First, each bilayer was equilibrated following the protocol recommended by CHARMM-GUI.[S2–S4] The equilibrated bilayer was then placed in a larger simulation box by extending one of the lateral membrane dimensions. Peptides were then placed near both membrane rims, after which the system was resolvated with water and ions.

Subsequently, the system underwent energy minimization, followed by a series of equilibration runs. The first equilibration was a short 20 ps $NVT$ run with a 1 fs time step. During this run, position restraints in all three spatial directions were applied to the Martini $PO_4$ beads (with a force constant $k_{pr} = 100$ kJ $\cdot$ mol$^{-1}$ $\cdot$ nm$^{-2}$) and peptide backbone beads ($k_{pr} = 1000$ kJ $\cdot$ mol$^{-1}$ $\cdot$ nm$^{-2}$). Next, a 100 ps $NPT$ run with a 2 fs time step and the same restraints ($k_{pr} = 50$ kJ $\cdot$ mol$^{-1}$ $\cdot$ nm$^{-2}$ on $PO_4$ beads; $k_{pr} = 1000$ kJ $\cdot$ mol$^{-1}$ $\cdot$ nm$^{-2}$ on peptide backbone beads) was conducted using an anisotropic barostat. This barostat acted independently only along the membrane's lateral dimensions, *i.e.*, along the extended dimension and parallel to the pore rim.

Following this equilibration, the position restraints were replaced with flat-bottom restraints acting on the phosphate beads to prevent rotation of the lipid stripe, as described in our previous work.[S1] Flat-bottom restraints were also applied to the peptide backbone beads to prevent the peptides from entering the bilayer region, moving to the opposite rim, or partially unbinding above or below the stripe, Figure S1. Peptides remained free to unbind from the rim into the pore region. With these restraints applied, a 200 ps $NVT$ simulation with a 2 fs time step was performed, leading to the formation or initial development of the lipid stripe while the peptides began to "cover" its rims. To generate starting configurations for umbrella sampling (US), the lipid stripe was then compressed along the pore-rim dimension during an $NPT$ simulation lasting up to 1 ns with a 5 fs time step. In this simulation, the box length along the $z$ direction was restrained using the PLUMED `RESTRAINT` action, with an equilibrium value of 5 nm and a force constant of $k_r = 1000\ \mathrm{kJ} \cdot \mathrm{mol}^{-1} \cdot \mathrm{nm}^{-2}$. This restraint compressed the simulation box because its initial length along the $z$ direction was approximately 7.5 nm. All equilibration simulations were carried out using the v-rescale thermostat[S5] with a coupling time of 1 ps and the Berendsen barostat[S6] with a coupling time of 1 ps for the initial $NPT$ simulation and 5 ps for the subsequent compression simulation.

From the compression simulation, lipid-stripe configurations with pore-rim sizes of 6.1, 6.2, and 6.3 nm were extracted. Each US window was first equilibrated for 4 ns in the $NVT$ ensemble and then for another 4 ns in the $NPT$ ensemble, using a 20 fs time step. The v-rescale thermostat[S5] was used in both simulations with a coupling time of 1 ps, while the Berendsen barostat[S6] was used during the $NPT$ simulation with a coupling time of 5 ps. After equilibration, the production runs were conducted as detailed in the main text. The flat-bottom restraints described above, together with the box-size restraint controlling the CV, were also applied in the US simulations, as illustrated in Figure S1. All other details of the simulation protocols for equilibration simulations followed the force field recommendations and were consistent with those used in the production runs.

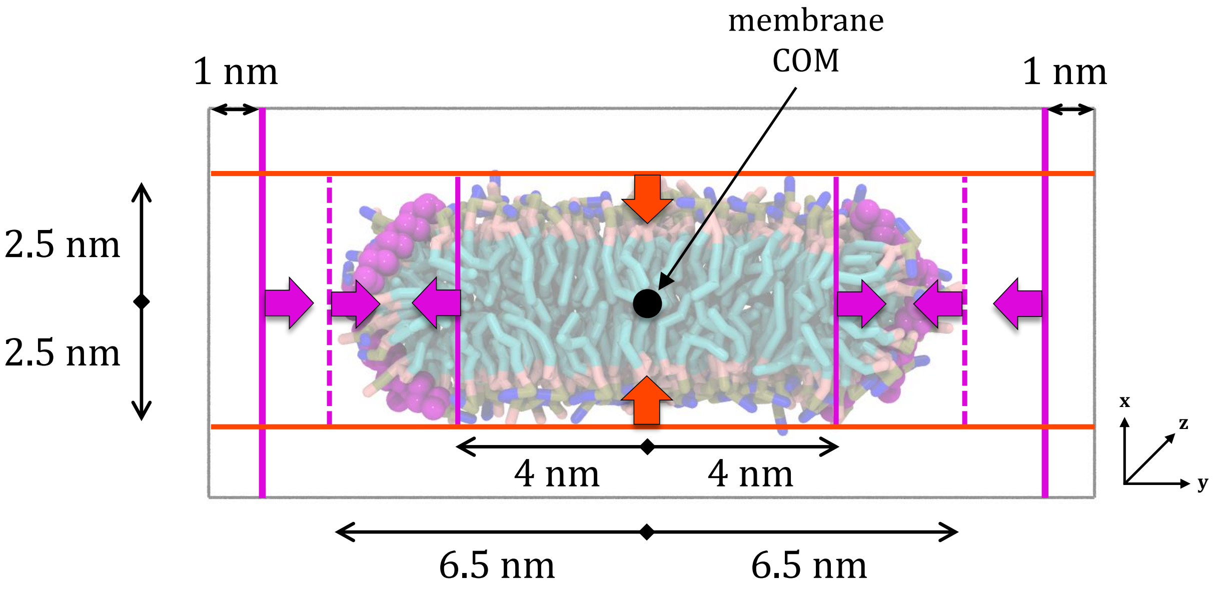


**Figure S1: Flat-bottom restraints used in simulations for line-tension estimation.** Lines indicate the positions of the restraint boundaries, while arrows indicate the directions of the forces acting on restrained beads. Purple denotes restraints acting on peptide backbone beads, whereas red denotes restraints acting on both peptide backbone and lipid phosphate beads. Dashed lines indicate restraints applied only during equilibration and removed for the production simulations. During the equilibration stages, the restraints located at 6.5 nm and 4 nm from the membrane center of mass along the $y$-axis had different values ([5.5, 6.0, 6.5] nm and [3.5, 4.0] nm, respectively), to first place the peptides at the membrane rim before the later equilibration stages and production simulations. An example peptide setup covering all equilibration stages, including the restraint positions and force constants used, is available in the Zenodo repository (the link will be made available following peer review).

### Martini Hybrid Model

In this work, we used a hybrid Martini 2.2 model that retains the Martini 2.2P representation of amino acids[S7] and polarizable water,[S8] while using a rescaled interaction matrix for peptide–peptide interactions[S9] and the standard Martini 2.2 interaction matrix[S7] for most other nonbonded interactions. This choice was motivated by three considerations. First, the polarizable version of Martini was required to accurately reproduce membrane line-tension trends.[S1] Second, the latest Martini model (version 3)[S10] has previously been reported to be less accurate than Martini 2 in capturing peptide sensing of membrane curvature,[S11] which is relevant here because the peptides localize at the curved pore rim. Third, rescaling peptide–peptide interactions was necessary to avoid excessive peptide aggregation, *e.g.*, at the membrane pore rim, and to obtain a more realistic description of peptide behavior. Moreover, scaling down peptide–peptide interactions also helps to emphasize the contribution of peptide–lipid interactions to line tension by reducing the tendency of peptides to associate strongly with each other. This choice is reasonable for toroidal pores, where peptides are expected to remain relatively loosely organized, in contrast to barrel-stave pores, in which specific peptide–peptide interactions are important for pore stability.[S12,S13]

Overestimation of peptide–peptide (protein–protein) interaction strength is a known issue of the Martini 2 model.[S9,S14] Our brief tests using two simple amphipathic peptides (the so-called LS4 and LS11 peptides,[S11] which contain only leucine and serine amino acids and have a clear amphipathic structure with different sizes of the polar patch) showed that the use of polarizable water does not resolve this issue and, in fact, instead increases the peptide–peptide interaction free energy in aqueous solution compared with the standard Martini 2 model, Table S1. This observation is consistent with previous reports indicating that polarizable water alone does not correct the overly strong protein–protein interactions in Martini 2.[S14–S16] Several scaling factors have been proposed to address this problem, based on either interaction of membrane proteins[S14] or soluble proteins.[S9] Here, we used a scaling factor of 30%, as suggested in the latter study,[S9] because the systems considered there are more representative of the peptides studied here.

The polarizable Martini 2.2P model modifies the interaction matrix for charged beads compared with the standard Martini 2.2 model.[S8] This raises the question of how these modifications should be combined with the rescaled peptide–peptide interaction matrix,[S9] and how charged beads belonging to peptides, lipids, and ions should be treated. For charged bead–water interactions, we retained the interactions from the original polarizable Martini parameterization,[S7,S8] thereby preserving its treatment of charged-bead hydration. For peptide–peptide interactions, however, we used the rescaled matrix[S9] without the charged-bead modifications introduced in the polarizable Martini 2.2P model.[S8] For peptide–lipid and lipid–lipid interactions, as well as the interactions of ions with peptides and lipids, we also retained the standard Martini 2.2 interaction matrix,[S7] without the polarizable-model modifications,[S8] see below for the rationale for this choice. Polarizable Martini 2 water was used throughout.[S8] We refer to this force field as Martini 2.2 hybrid polarizable and scaled (M2.2hps).

This choice was motivated in part by the variable performance of polarizable Martini across different systems and observables and by the comparatively limited benchmarking available for the peptide–lipid and lipid–lipid properties relevant to the present study.[S7,S8,S17,S18] We therefore performed two tests of our own to assess the suitability of the selected hybrid model. First, polarizable water combined with the standard Martini 2.2 interaction matrix was sufficient to reproduce the experimentally consistent[S19] line-tension trend observed in our previous work using the original Martini 2.2P model,[S1] namely, the lower line tension of POPG compared with POPC, Table S2. Second, for pure lipid bilayers, combining polarizable water with the standard Martini 2.2 interaction matrix produced area-per-lipid values closer to the AA-MD results than when the charged-bead modifications of the polarizable model were applied, Table S3. This trend was observed not only for the POPE:POPG 3:1 composition used in this work, but also for POPC. We consider this particularly relevant because the present study focuses on lipid organization at the pore rim and on peptidemediated shielding of hydrophobic lipid tails from water, making lipid packing an important property of the model.

**Table S1:** Free energy of interaction (in $kJ \cdot mol^{-1}$) between two peptides in aqueous solution using different versions of the Martini 2 force field. Reported values correspond to the free-energy minima obtained using the accelerated weight histogram method[S20] implemented in GROMACS.[S21] The collective variable was the distance between the centers of mass of the two peptides, either two LS4 or two LS11 peptides. Values were calculated from three independent 1 $\mu$s replicas, with the corresponding standard deviations reported.

| Peptide | Force field | | |
|---|---|---|---|
| | M2.2 | M2.2 scaled (10% scaling[S14]) | M2.2 polarizable |
| `LSLLLLLLSLLLSLLLLLLSL` (LS4) | –60.9 ± 2.6 | –55.5 ± 3.1 | –67.1 ± 2.3 |
| `LSSLLSLLSSLLSSLSSLLSS` (LS11) | –47.5 ± 5.0 | –42.6 ± 1.0 | –53.5 ± 3.0 |

**Table S2:** Line-tension predictions (in pN) for peptide-free systems obtained using the Rapid method with different versions of the Martini 2 force field. The same simulation setup as in our original work was used.[S1]

| Lipid composition | Force field | | |
|---|---|---|---|
| | M2.2[a] | M2.2 polarizable[a] | **M2.2hps (this work)** |
| POPC | 57.3 ± 0.0 | 49.2 ± 0.6 | **56.1 ± 1.0** |
| POPE | 76.2 ± 1.0 | 55.7 ± 0.6 | **67.7 ± 0.5** |
| POPE:POPG 3:1 | – | 54.1 ± 0.1 | **64.4 ± 0.6** |
| POPG | 65.1 ± 0.3 | 38.8 ± 0.6 | **49.8 ± 0.4** |

[a] Values taken from Ref. S1.

**Table S3:** Area per lipid (in $nm^2$) for lipid bilayers obtained from CG-MD simulations using different versions of the Martini 2 force field, compared with AA-MD simulations using the CHARMM36 force field.[S22]

| Lipid composition | Force field | | | |
|---|---|---|---|---|
| | M2.2 | M2.2 polarizable | **M2.2hps (this work)** | CHARMM36 |
| POPE:POPG 3:1 | 0.619 | 0.643 | **0.630** | 0.590 |
| POPC | 0.656 | 0.656 | **0.650** | 0.645 |

#### Application of Rapid CV

Our original setup with Rapid CV used 21 umbrella sampling (US) windows to estimate the line tension of peptide-free systems.[S1] To make this approach feasible for use with Evo-MD-FE, we reduced the setup to only 3 US windows, using a weaker force constant over a narrower range of box sizes. Note that the system used here, and therefore the pore size across the periodic boundaries, was slightly larger than in our previous work[S1] to ensure that peptide–peptide interactions across the pore were prevented. The setup used here contained more water beads (6,000 vs. 5,000) and therefore more ion pairs to maintain a comparable physiological salt concentration (65 vs. 54).

Comparison of the setups, Table S4, showed that 3 US windows were sufficient to adequately reproduce the line-tension values for peptide-free systems. In peptide-containing systems, larger deviations can be expected. However, as discussed for the Evo-MD-FE in the main text, full convergence of the absolute line-tension value is not required for the evolutionary search, which relies primarily on the relative ranking of candidate sequences. The reduced setup was therefore used to provide a consistent comparative fitness metric for peptide ranking rather than fully converged absolute line-tension values for every peptide.

The US histogram counts from simulations of a representative system containing the LT1 peptide are shown in Figure S2. As can be seen, there is substantial overlap between the three simulated windows. Sampling at larger box sizes is somewhat lower because, despite equilibrium distances of 6.1, 6.2, and 6.3 nm, the system naturally drifted toward smaller values under the relatively weak restraint. To verify that this did not affect the analysis, we tested whether changing the linear-fit range to 6.05–6.25 nm meaningfully alters the resulting line-tension estimate. Figure S3 shows excellent agreement between the two fitting ranges for the data from run #1 (Table S5), with differences comparable to those expected between independent replicas analyzed using the same fitting range. The only outlier corresponds to an increase in line tension, *i.e.*, to a state in which peptide binding is unfavorable; in this case, the free-energy profile is less well sampled and more sensitive to the exact region chosen for the fit, but this peptide is of limited relevance to the present search.

**Table S4:** Line-tension predictions (in pN) for peptide-free systems obtained using the Rapid method with different numbers of US windows using the M2.2hps force field. For simulations with 21 US windows, the setup from our previous work was used.[S1] For simulations with 3 US windows, two independent replicas were performed, and the mean value with the corresponding standard deviation is reported.

| Lipid composition | 21 US × 150 ns original setup[S1] 6.0–6.6 nm | 21 US × 150 ns this work's setup 6.0–6.6 nm | 3 US × 300 ns this work's setup 6.1–6.3 nm |
|---|---|---|---|
| POPE:POPG 3:1 | 64.4 ± 0.6[a] | 65.8 ± 0.0 | 62.0 ± 0.7 |

[a] This value was used as the reference line tension throughout this work.

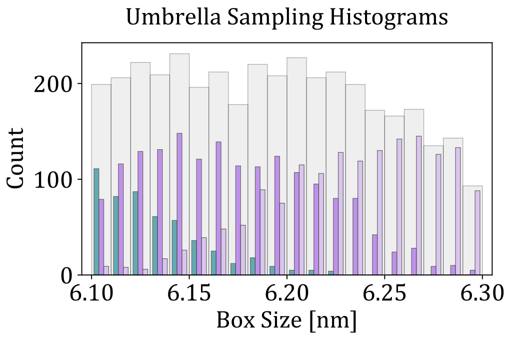


**Figure S2:** Umbrella-sampling histograms from simulations of a representative system containing the LT1 peptide using the reduced 3 US-window setup. Colored histograms show the sampled box-size distributions of the three individual US windows, while the gray histogram shows their sum.

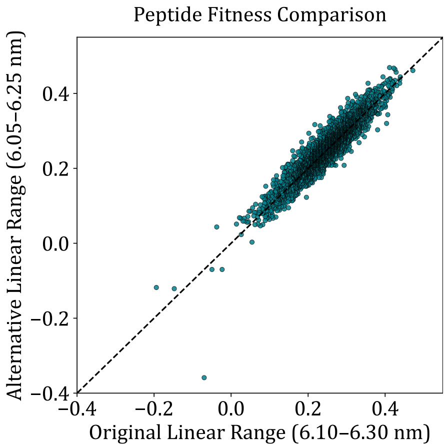


**Figure S3:** Comparison of peptide fitness values obtained using different linear fitting ranges for the free energy profiles. Peptide fitness calculated using the 6.10–6.30 nm fitting range (used in this work) is compared with values obtained using the alternative 6.05–6.25 nm range. Each point represents one set of US simulations. The dashed line is the identity line.

### Evo-MD-FE Optimization Runs

#### Generation of Children in the "Face" Method

In the "Face" method of children generation, each parent peptide sequence was mapped onto a helical wheel and aligned according to its hydrophobic moment. The helical wheel was then divided into two faces of equal size along an axis perpendicular to the hydrophobic moment, and the corresponding faces were exchanged between two parent peptides. Each pair of parent peptides generated two child peptides; if adding both children would exceed the prescribed population size for the iteration, only one child was added.

To convert the resulting helical-wheel representations back into primary sequences, one parent was used as the positional reference for each child. Residues retained from this parent remained at their original primary-sequence positions, whereas residues from the corresponding helical face of the second parent were placed at the primary-sequence positions of the reference parent corresponding to their locations on the helical face. Thus, the transferred residues preserved their location on the helical face, but not necessarily their original positions in the primary sequence of the second parent. The second child was generated analogously, using the other parent as the positional reference.

#### Performed Optimization Runs

**Table S5:** Settings of the Evo-MD-FE optimization runs performed in this work.

| # | Scheme | AA set | Mut. rate | # Iter. / Pep./iter. / Total peps. |
|---|---|---|---|---|
| 1 (main text) | Face | 1/18 | ALMKEQSFWY | 30 / 128 / 3,723 |
| 2 (replicate of #1) | Face | 1/18 | ALMKEQSFWY | 20 / 128 / 2,486 |
| 3 | Face | $3/18 \rightarrow 1/18$ ($-0.1/18$/iter.) | AGLVCMKREDQNSTFWYH | 36 / 128 / 4,976 |
| 4 | Block | 1/18 | ALMKEQSFWY | 20 / 128 / 2,284 |

#### Additional Analysis of the Run #1 Discussed in the Main Text

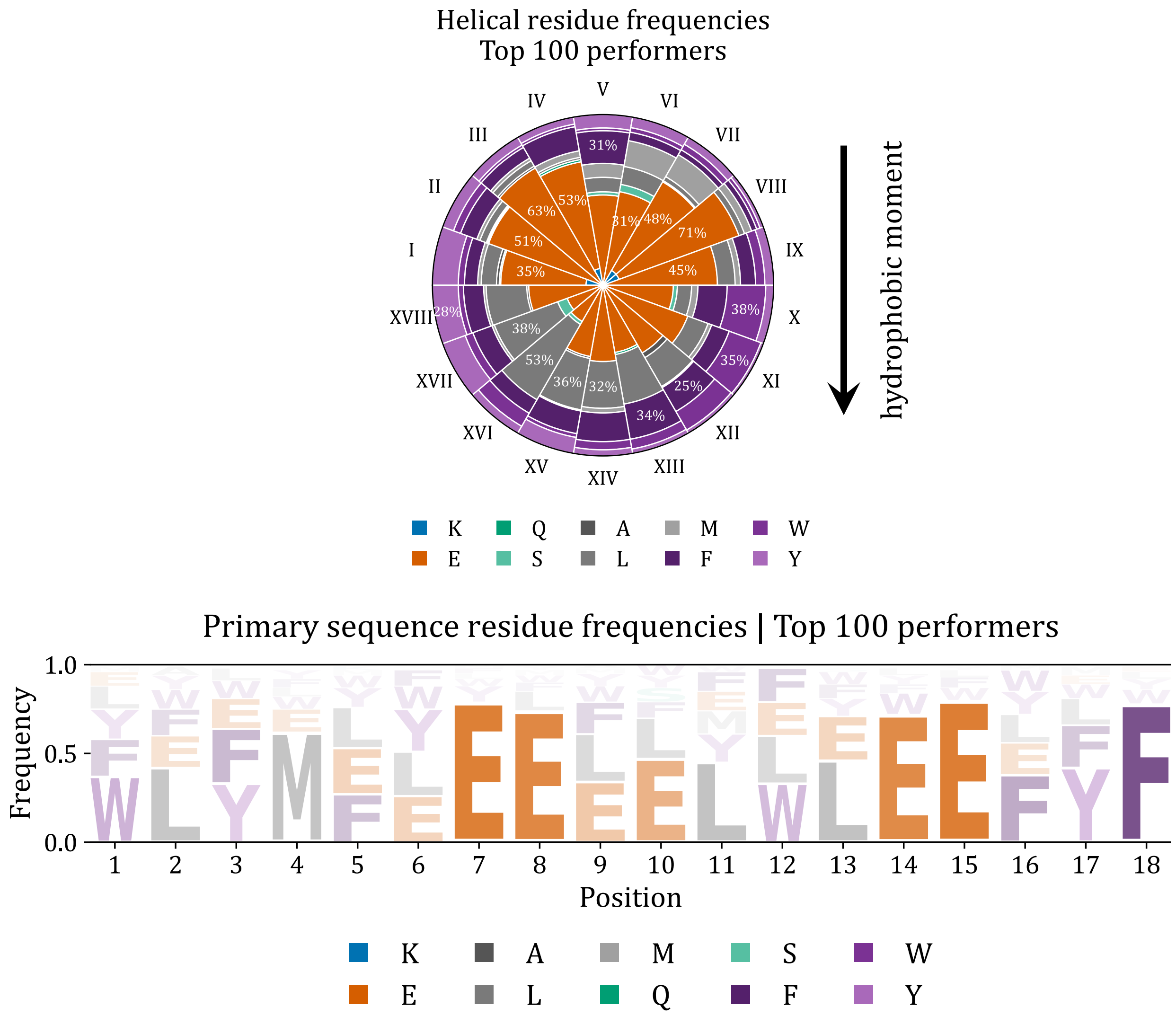


**Figure S4:** Residue frequencies in the helical wheel (top) and in the primary sequence (bottom). All definitions, plotting conventions, and analysis details are identical to those described for Figure 2 in the main text.

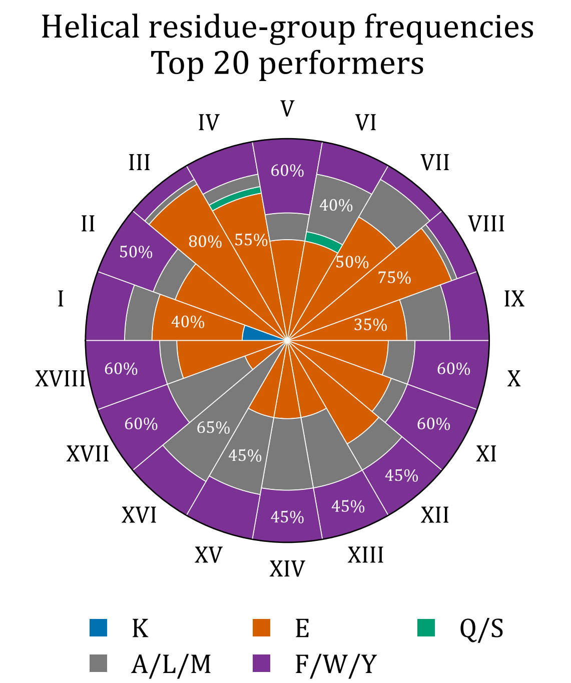


**Figure S5:** Residue-group frequencies in the helical wheel for the top 20 performers. All definitions, plotting conventions, and analysis details are identical to those described for Figure 2 in the main text.

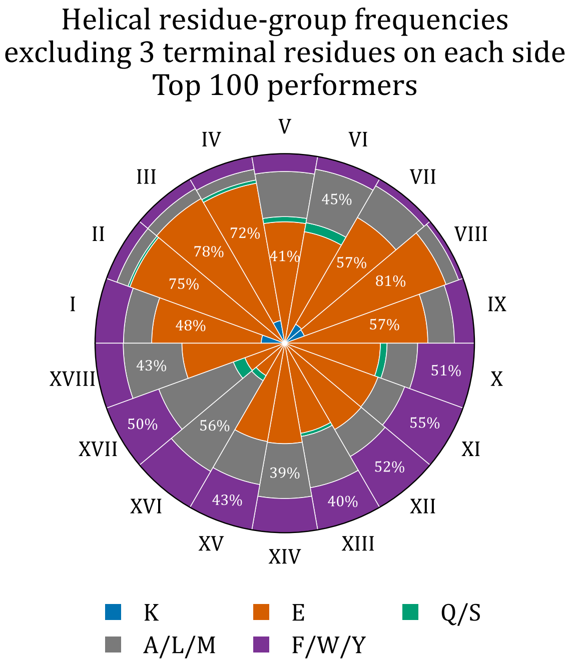


**Figure S6:** Residue-group frequencies in the helical wheel calculated after excluding the first and last three residues of each primary sequence. All definitions, plotting conventions, and analysis details are identical to those described for Figure 2 in the main text.

### Results from the Additional Runs

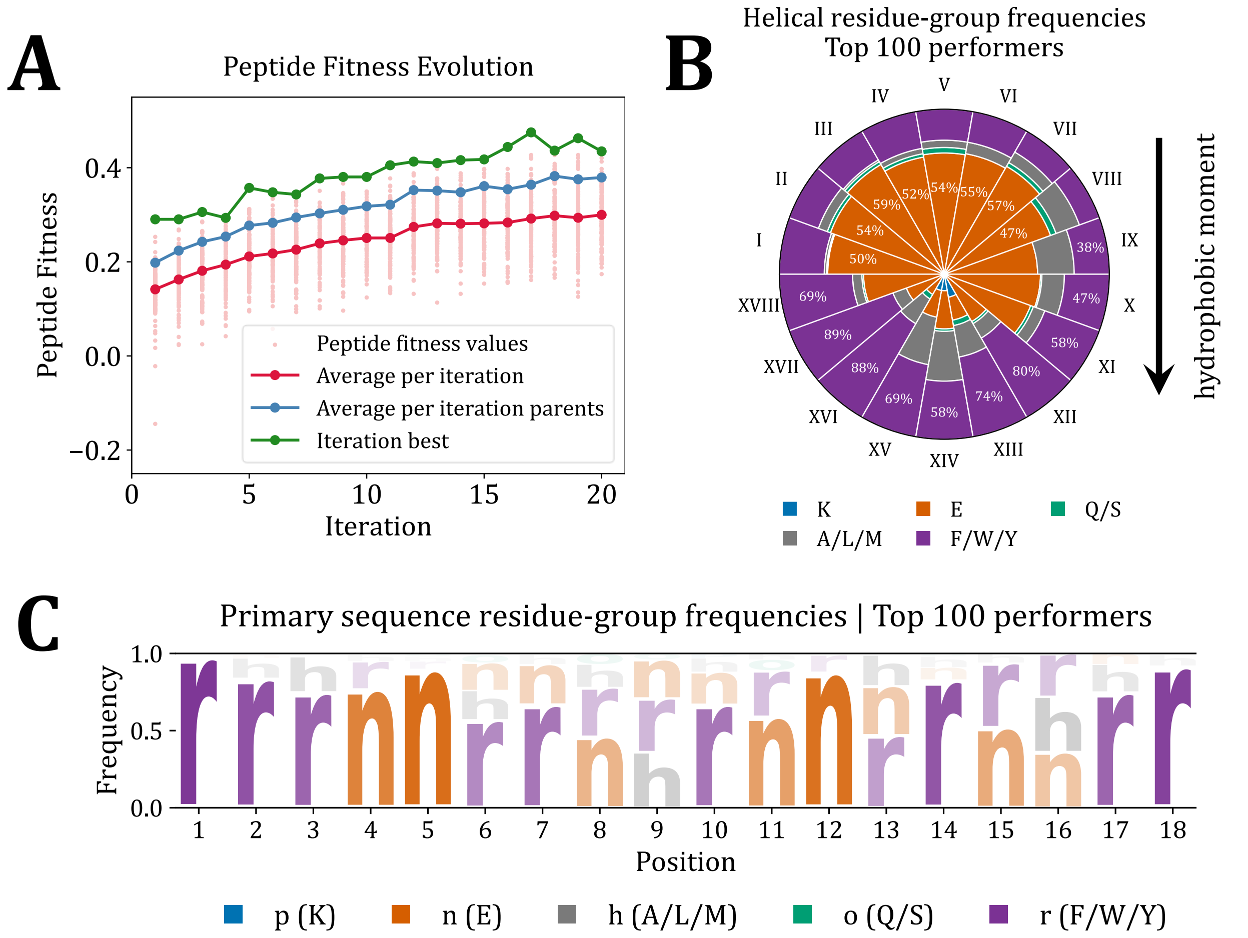


**Figure S7: A–C) Evo-MD-FE optimization run #2**, see Table S5. A) Evolution of peptide fitness. B) Residue-group frequencies in the helical wheel. C) Residue-group frequencies in the primary sequence. All definitions, plotting conventions, and analysis details are identical to those described for Figure 2 in the main text.

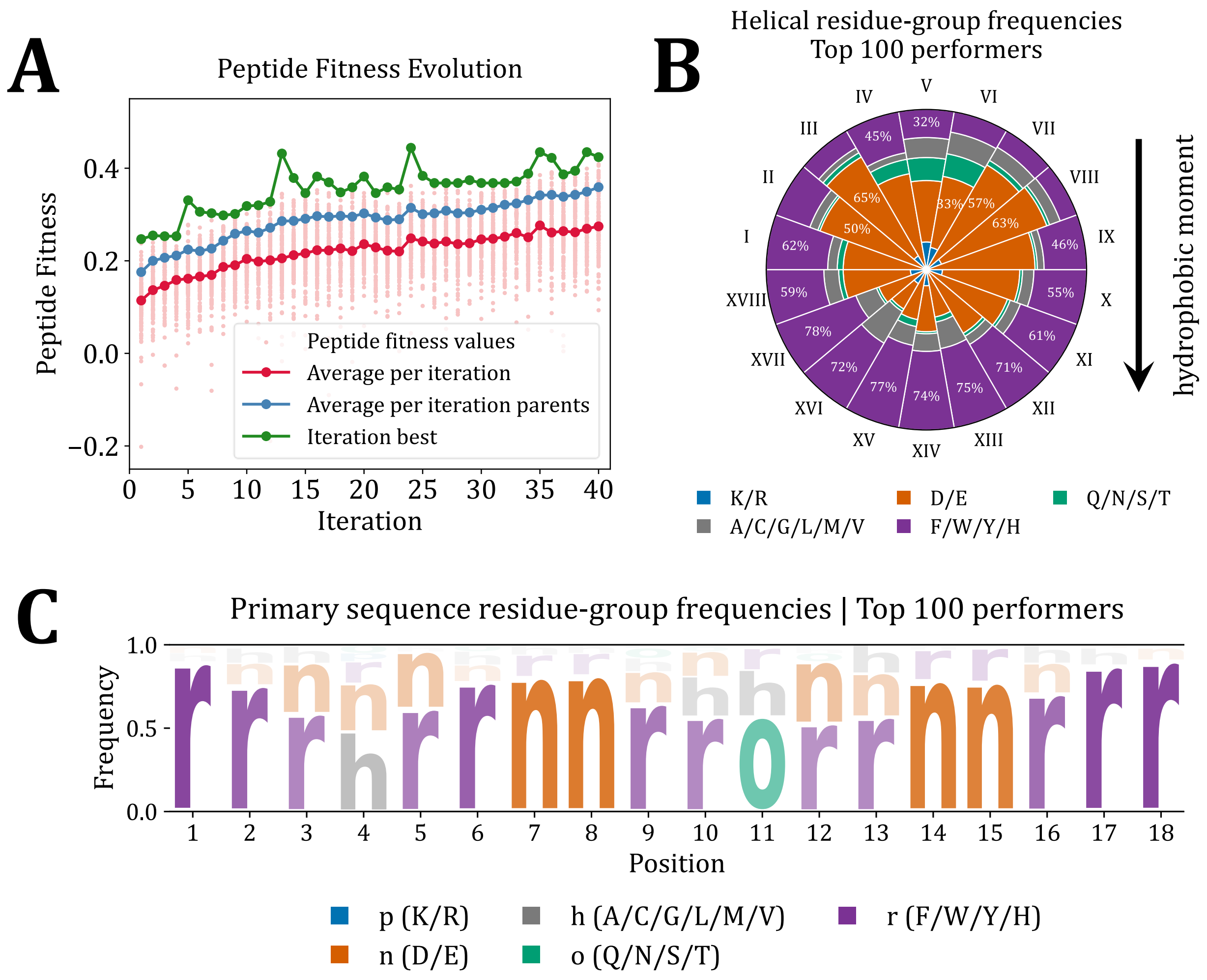


**Figure S8:** **A–C) Evo-MD-FE optimization run #3**, see Table S5. A) Evolution of peptide fitness. B) Residue-group frequencies in the helical wheel. C) Residue-group frequencies in the primary sequence. All definitions, plotting conventions, and analysis details are identical to those described for Figure 2 in the main text.

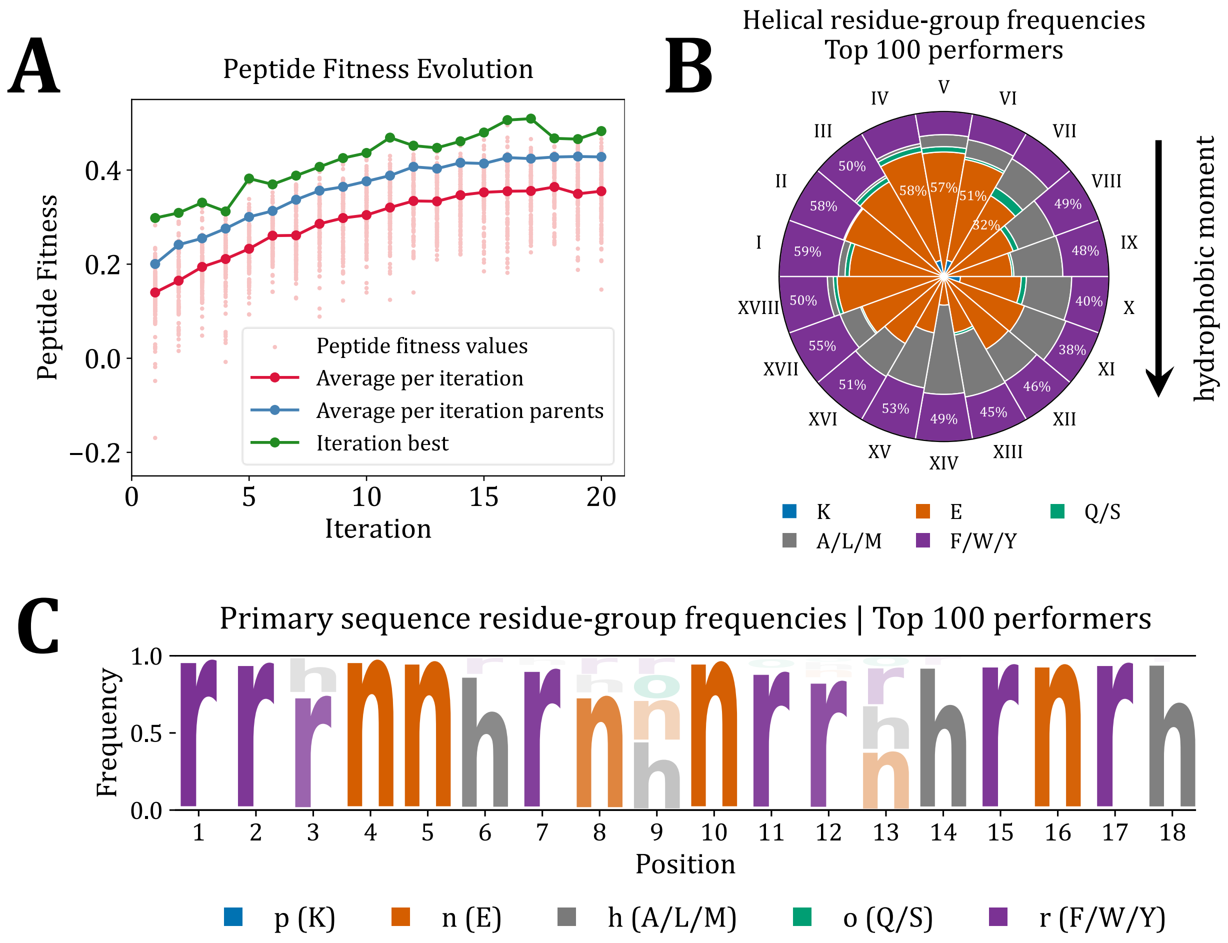


**Figure S9:** **A–C) Evo-MD-FE optimization run #4**, see Table S5. A) Evolution of peptide fitness. B) Residue-group frequencies in the helical wheel. C) Residue-group frequencies in the primary sequence. All definitions, plotting conventions, and analysis details are identical to those described for Figure 2 in the main text.

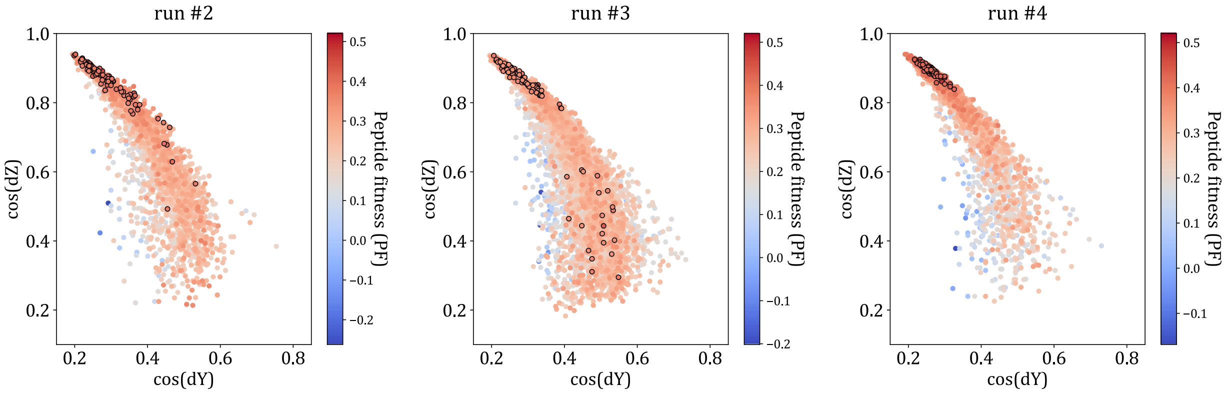


**Figure S10: Peptide orientation at the pore rim** from additional Evo-MD-FE runs, see Table S5. All definitions, plotting conventions, and analysis details are identical to those described for Figure 3 in the main text.

### Supplementary Data and Discussion

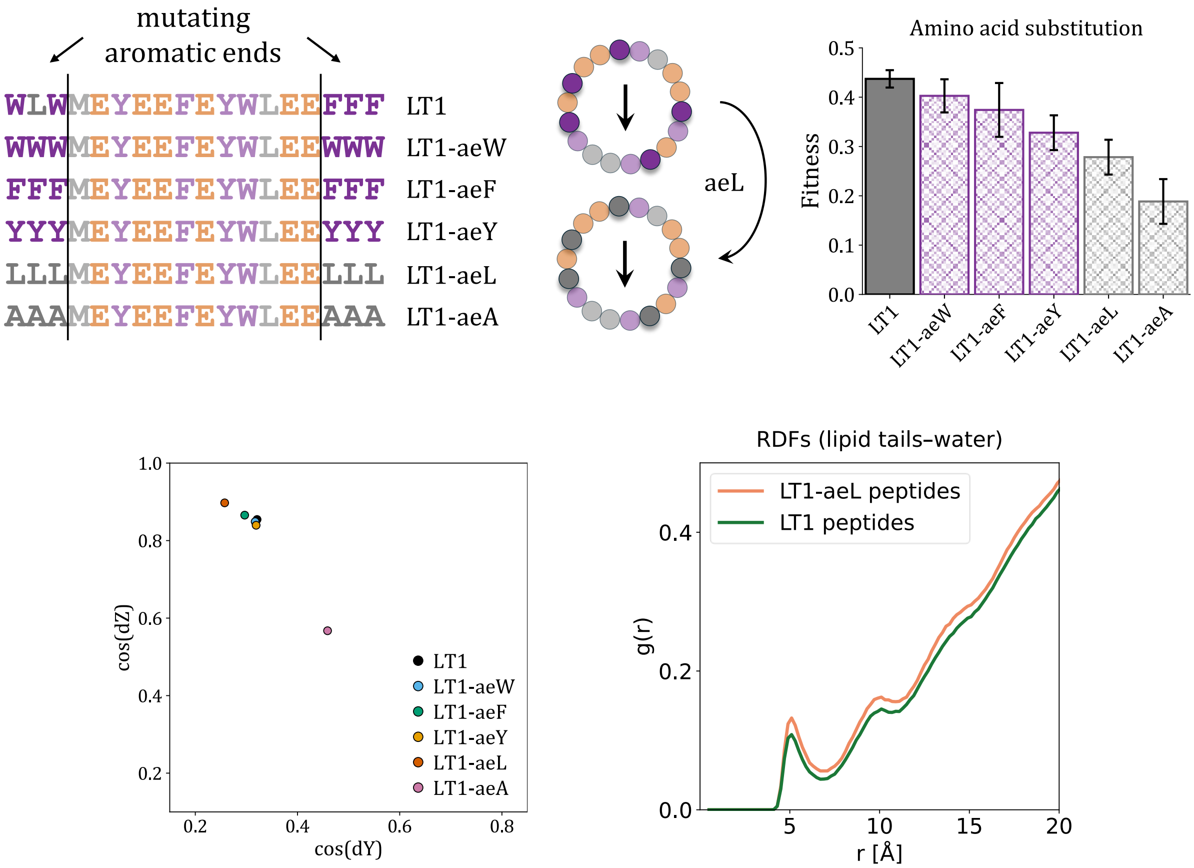


**Figure S11: Mutating aromatic ends.** The first and the last three residues in LT1 were mutated to either W, F, Y, L, or A, while the remainder of the sequence was retained. The fitness values of LT1 and the resulting variants are compared (top). The error bars represent the standard deviation across replicate simulations. Their orientations at the pore rim are compared in the bottom-left panel. RDFs, calculated as in Figure 3, are shown for LT1 and LT1-aeL (bottom-right) to illustrate the reduced shielding of lipid tails from water by LT1-aeL peptides, which is consistent with their poorer performance despite similar orientation and localization at the pore rim.

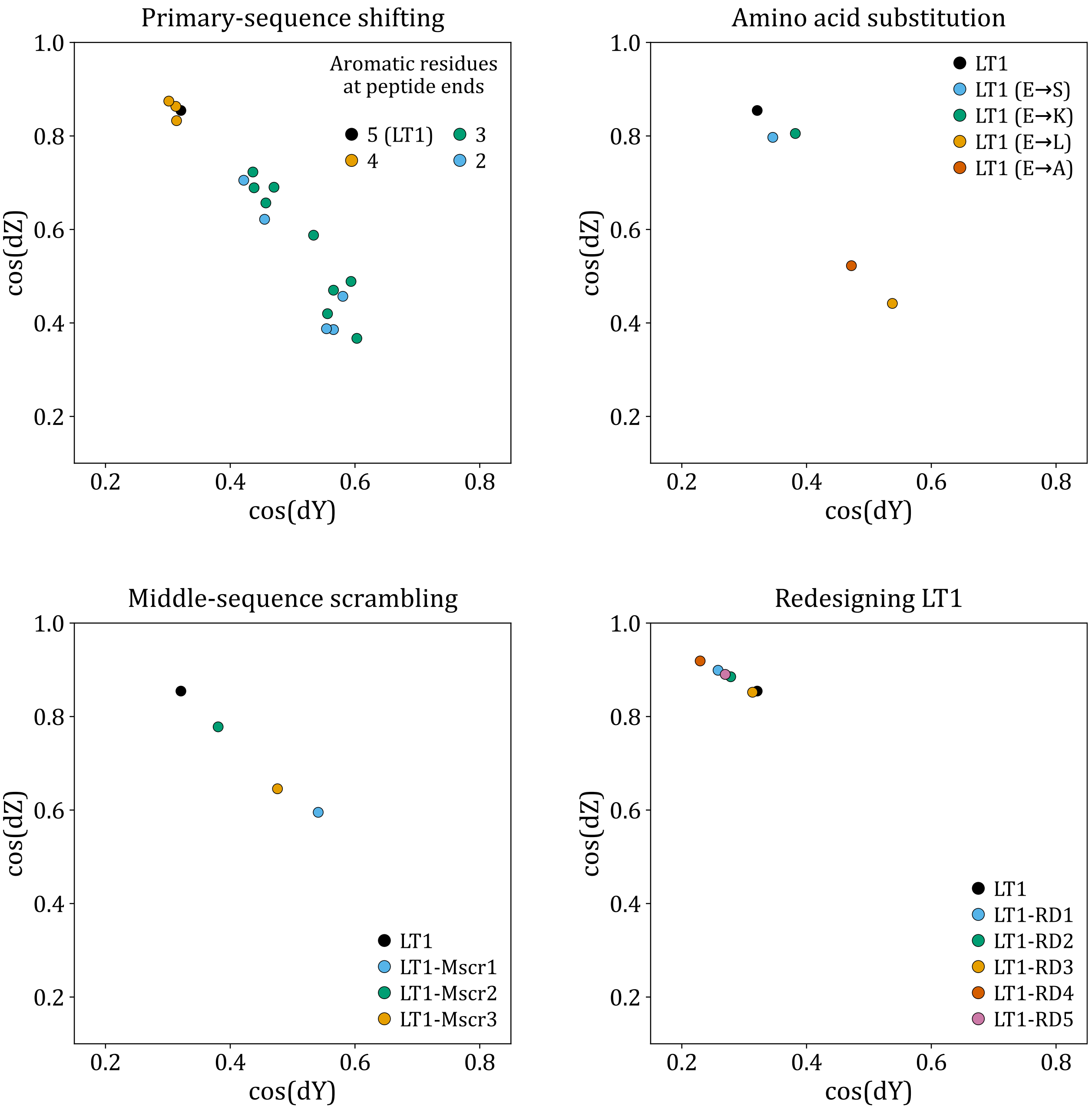


**Figure S12: Peptide orientation at the pore rim** upon the sequence modifications shown in Figure 4.

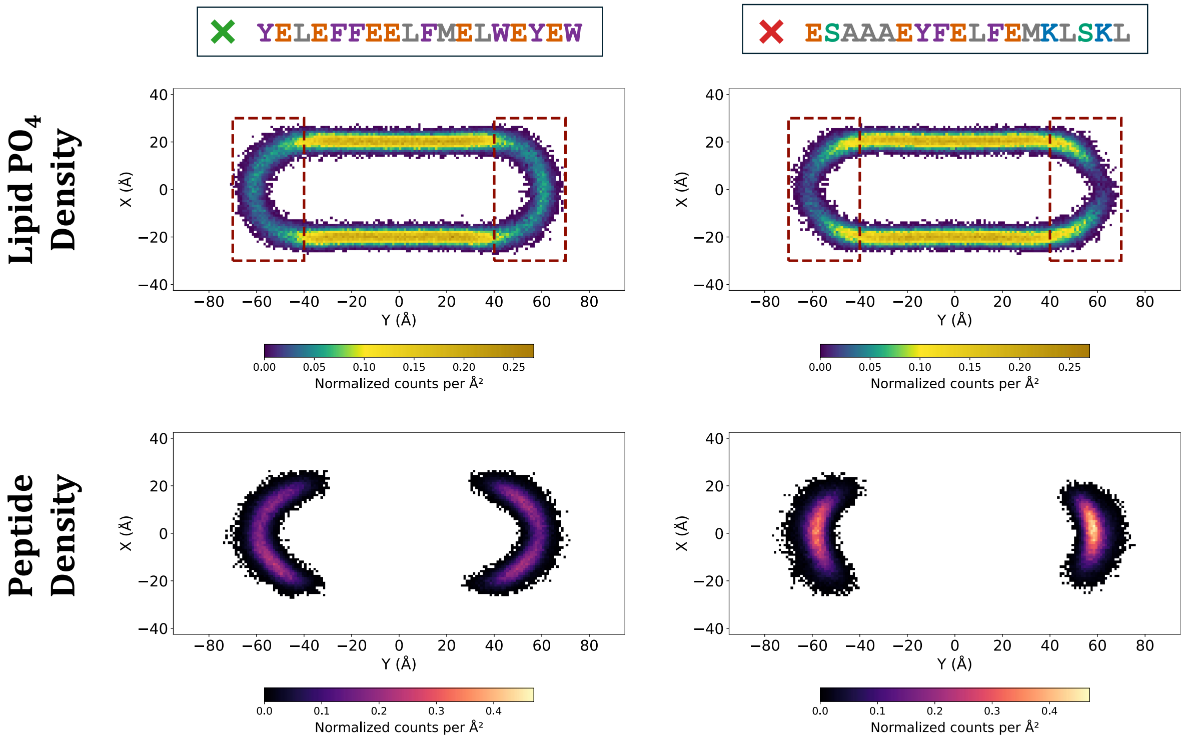


**Figure S13:** Two-dimensional density maps of the stripe cross-sections from CG-MD simulations for the peptide “outliers” shown in Figure 5 and discussed in the main text.
